# Gut regulates brain synaptic assembly through neuroendocrine signaling pathway

**DOI:** 10.1101/2021.01.29.428811

**Authors:** Yanjun Shi, Lu Qin, Zhiyong Shao

**Affiliations:** Department of Neurosurgery, State Key Laboratory of Medical Neurobiology and MOE Frontiers Center for Brain Science, Institutes of Brain Science, Zhongshan Hospital, Fudan University, Shanghai 200032, China

**Author notes:** Corresponding author and lead contact Electronic address.

**Keywords:** Gut-brain axis, Synaptic development, Wnt, CWN-2, Endocrine, Neuropeptide/NLP-40, GPCR/AEX-2

## Abstract

The gut-brain axis plays an essential role in regulating neural development in response to environmental stimuli, such as microbes or nutrients. Defects in gut-brain communication can lead to various neurological disorders. However, it remains unknown whether gut plays any intrinsic role in regulating neuronal development. Through a genetic screen in *C. elegans*, we uncovered that an intrinsic Wnt-endocrine pathway in gut regulates synaptic development and neuronal activity in brain. Specifically, the Wnt signaling upregulates the expression of the neuropeptide NLP-40 in the gut, which then facilitates presynaptic assembly through the neuronal expressed GPCR AEX-2 receptor during development. The NLP-40 acts most likely through modulating neuronal activity and promoting synaptic protein trafficking. Therefore, this study reveals a novel role of gut in synaptic development in the brain.

## INTRODUCTION

The gastrointestinal (GI) tract plays a critical role in regulating brain development and function. It not only provides the nutrient, but also senses the gut environmental stimuli. The gut-brain communication is recognized as gut-brain axis. Through this axis, diets and gut microbes regulate neurogenesis, microglia and astrocyte activation, myelination, blood brain barrier permeability, synaptic pruning and plasticity (Cryan et al., 2019). Dysregulation in the gut-brain axis is correlated with various neurodevelopmental or neurodegenerative disorders (Chidambaram et al., 2020; Kowalski and Mulak, 2019; Li and Zhou, 2016; Mulak and Bonaz, 2015; Quigley, 2017). Although accumulative evidence indicates that gut signaling affects neuronal development and function in the brain, it remains largely unknown if it plays any intrinsic role in brain development.

Gut-brain communication is mainly mediated through three pathways: afferent nervous system, immune system and endocrine system (Wang and Wang, 2016). Among those, the endocrine system is the most complex and highly conserved in both vertebrates and invertebrates (Bernhard Kleine and Rossmanith; Campbell et al., 2004; Dimaline and Dockray, 1994; Nassel et al., 2019). Upon stimulation by nutrients or gut microbiota, enteroendocrine cells (EECs) release neuropeptides or hormones such as cholecystokinin (CCK), ghrelin, leptin, peptide tyrosine-tyrosine (PYY), and glucagon-like peptide-1 (GLP-1) which can travel a long distance as endocrine factors through binding GPCR or other types of receptors to regulate the feeding behavior and the physiological function of the nervous system (Dockray, 2013; Wang et al., 2013). However, it is largely undetermined whether the intestinal endocrine system is involved in neurodevelopment or synaptic formation.

In this study, we uncovered that a canonical Wnt pathway in the intestine regulates synaptic development in the nerve ring, which is the nematode’s brain. We further demonstrated that the Wnt signaling functions through upregulating the neuroendocrine molecule NLP-40, which then promotes the synaptic assembly most likely through modulating the neuronal activity. Therefore, our study reveals a novel gut-brain interaction underlying synaptogenesis in brain.

## RESULTS

### Intestinal Wnt signaling regulates AIY presynaptic formation

*C. elegans* AIY neurons are a pair of bilateral symmetric interneurons located in the nerve ring, which is a structure analogous to vertebrate’s brain (Fig. 1A)(White et al., 1986). AIY neurites form presynapses with a stereotypic and highly reproducible pattern (Fig. 1A’)(White et al., 1986).

**Fig. 1.**
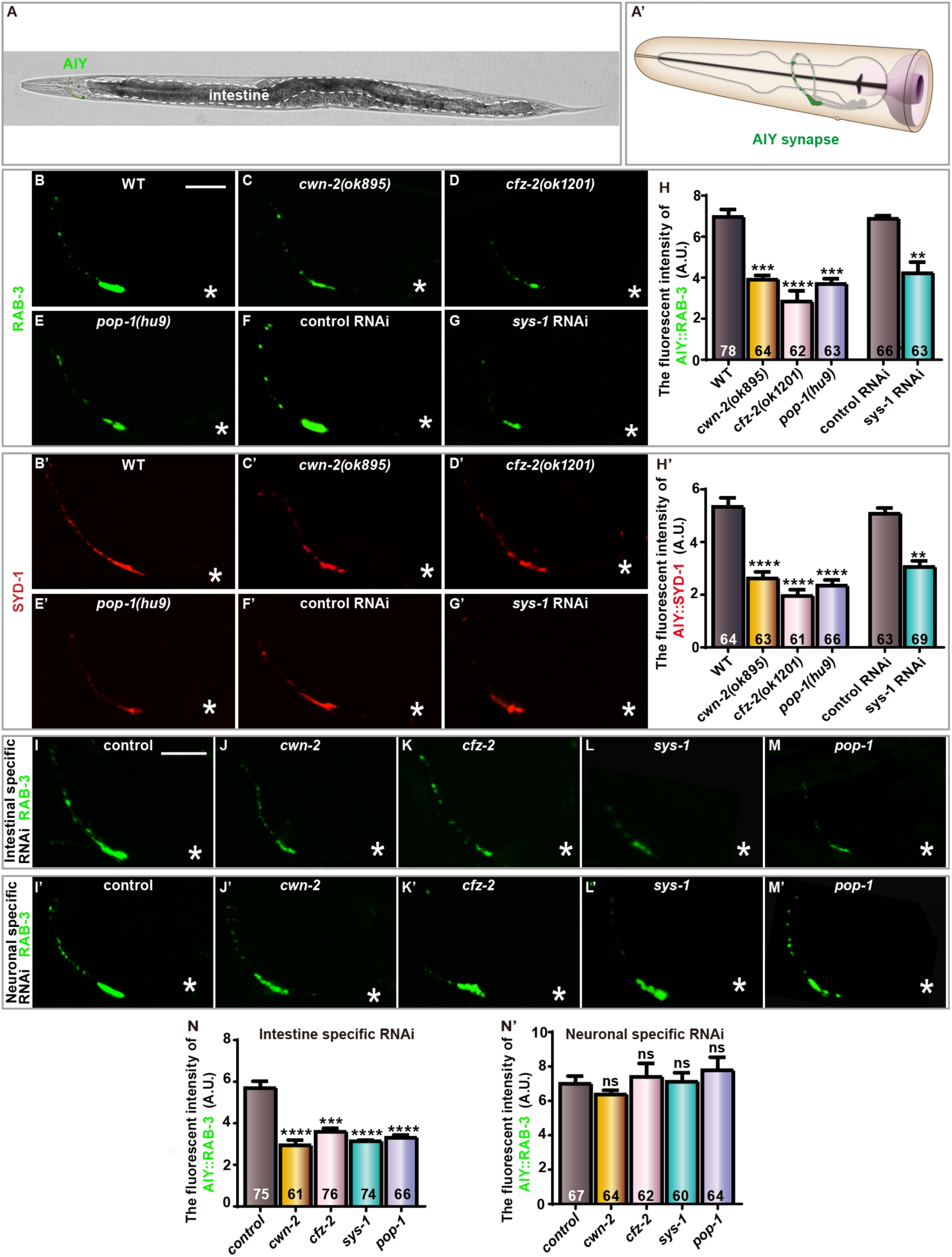
AIY presynaptic assembly requires Intestinal Wnt signaling. **(A)** A bright field image of a wild-type adult *C. elegans*; the position of the AIY interneurons in the head are indicated with green color, and the intestine is outlined with dashed lines. (A’) A cartoon diagram of the *C. elegans* head modified from wormatlas with permission. The AIY processes and soma are indicated in grey, and the presynaptic sites are marked with green color (Altun et al., 2002–2019). **(B-G’)** Confocal images of the AIY labeled with synaptic vesicle marker GFP::RAB-3(B-G, green) and synaptic active zone marker GFP::SYD-1(B’-G’, pseudo-red) in wild-type (B, B’), *cwn-2(ok895)* (C, C’), *cfz-2(ok1201)* (D, D’), *pop-1(hu9)* (E, E’), and *sys-1* RNAi (G, G’) animals. The scale bar in (B) is 10μm and applied to (C-G’). **(H-H’)** Quantification of fluorescent intensity of RAB-3 (H) and SYD-1 (H’) for the indicated genotypes. **(I-M’)** Confocal images of AIY labeled with mCherry::RAB-3 (pseudo-GFP) (I-M) or GFP::RAB-3 (I’-M’) of tissue-specific RNAi treatment for control (I-I’), *cwn-2*(J-J’), *cfz-2*(K-K’), *sys-1*(L-L’) and *pop-1*(M-M’). The scale bar in (I) is 10 μm and applied to (J-M’). **(N-N’)** Quantification of GFP::RAB-3 fluorescent intensity for the indicated RNAi treatment. For (H-H’, N-N’), data are averaged from at least three biological replicates. The total number (N) of independent animals assayed is indicated in the bars. Statistical analyses are based on one-way ANOVA followed by Dunnett’s test (for mutants) or two-tailed Student’s t-test (for RNAi). ns: not significance (p>0.05), **p<0.01, ***p<0.001, ****p<0.0001 as compared to the wild-type or control RNAi. Error bars represent SEM.

Wnt signaling pathways are highly conserved and play complicated roles in regulating synaptic formation, plasticity and maintenance (Dickins and Salinas, 2013; McLeod and Salinas, 2018; Yang and Zhang, 2020). We previously found that components in the canonical Wnt signaling pathway including Wnt/CWN-2, Frizzled/CFZ-2, Disheveled/DSH-2, β-catenin/SYS-1 and TCF/POP-1 are required for the AIY presynaptic morphogenesis (Shi et al., 2018). To address if the Wnt signaling pathway regulates the AIY presynaptic formation, we first quantified the fluorescent intensity of the AIY synaptic vesicle marker GFP::RAB-3 in *cwn-2(ok895)*, *cfz-2(ok1201), pop-1(hu9)* loss-of-function mutant and *sys-1* RNAi animals. We found that loss of this Wnt signaling resulted in a robust reduction of AIY synaptic marker GFP::RAB-3 (GFP intensity was reduced by 43.94%, 59.32%, 47.10% and 38.54% in *cwn-2(ok895)*, *cfz-2(ok1201), pop-1(hu9)* and *sys-1* RNAi. P<0.001, 0.0001 and 0.001 compared with WT for mutants, respectively, and P<0.01 for comparison between *sys-1* and control RNAi; Fig. 1B-H). We then examined whether the regulatory role of Wnt was specific to synaptic vesicle, so we examined the active zone marker GFP::SYD-1. Similar to GFP::RAB-3, GFP::SYD-1 was significantly reduced in these Wnt mutants (Fig. 1B’-H’). To exclude the possibility that the reduction of the presynaptic marker was specific to the GFP fluorophore, we quantified the mCherry::RAB-3 and obtained similar results (Supplementary Fig. 1A-G). These results collectively suggest that the canonical Wnt signaling is required for AIY presynaptic assembly.

To determine the site of action for the Wnt signaling, we knocked down the Wnt receptor and the downstream signaling components through neural or intestinal specific RNAi. Interestingly, we found robust reduction of AIY presynaptic markers after knocking down Wnt signaling in the intestine (the GFP intensity was reduced by 48.33%, 36.91%, 44.82% and 41.83% in *cwn-2, cfz-2, sys-1* and *pop-1* RNAi. P<0.0001, 0.001, 0.0001 and 0.0001 compared to the control RNAi, respectively; Fig. 1I-N), but not in the nervous system (Fig. 1I’-N’). Consistently, the *cfz-2(ok1201)* mutants could only be rescued by the intestine-specific, but not by the neuronal-specific, expression of *cfz-2* cDNA (Supplementary Fig. 2D-F). These data collectively suggest that CWN-2 acts through canonical Wnt components CFZ-2/SYS-1/POP-1 in the gut to regulate AIY presynaptic formation.

### Intestinal neuropeptide/NLP-40 is required for synaptogenesis

Gut can communicate with brain through neuroendocrine signaling pathway (Campbell et al., 2004; Dimaline and Dockray, 1994; Nassel et al., 2019). To test whether Wnt signaling in the gut regulates AIY synaptogenesis through neuroendocrine or neuropeptide signaling, we first examined if EGL-3, a key proprotein convertase (PC2) involved in neuropeptide maturation (Li and Kim, 2008), was required for the AIY presynaptic assembly. Indeed, the fluorescence intensity of AIY presynaptic markers was robustly reduced in *egl-3(n589)* mutants (P<0.01, Fig. 2A), suggesting that neuropeptides are required for the AIY presynaptic assembly. There are 115 neuropeptide encoding genes identified in *C. elegans* genome (Li and Kim, 2008; Nathoo et al., 2001). We screened most neuropeptide genes enriched in the intestine through RNAi for AIY synaptic regulators (Fig. 2B)(Li and Kim, 2008; Pauli et al., 2006; Spencer et al., 2011; Wang et al., 2013). Interestingly, *nlp-40* knockdown displayed a dramatic reduction of the synaptic marker, suggesting that it is required for the AIY presynaptic assembly (Fig. 2C).

**Fig. 2.**
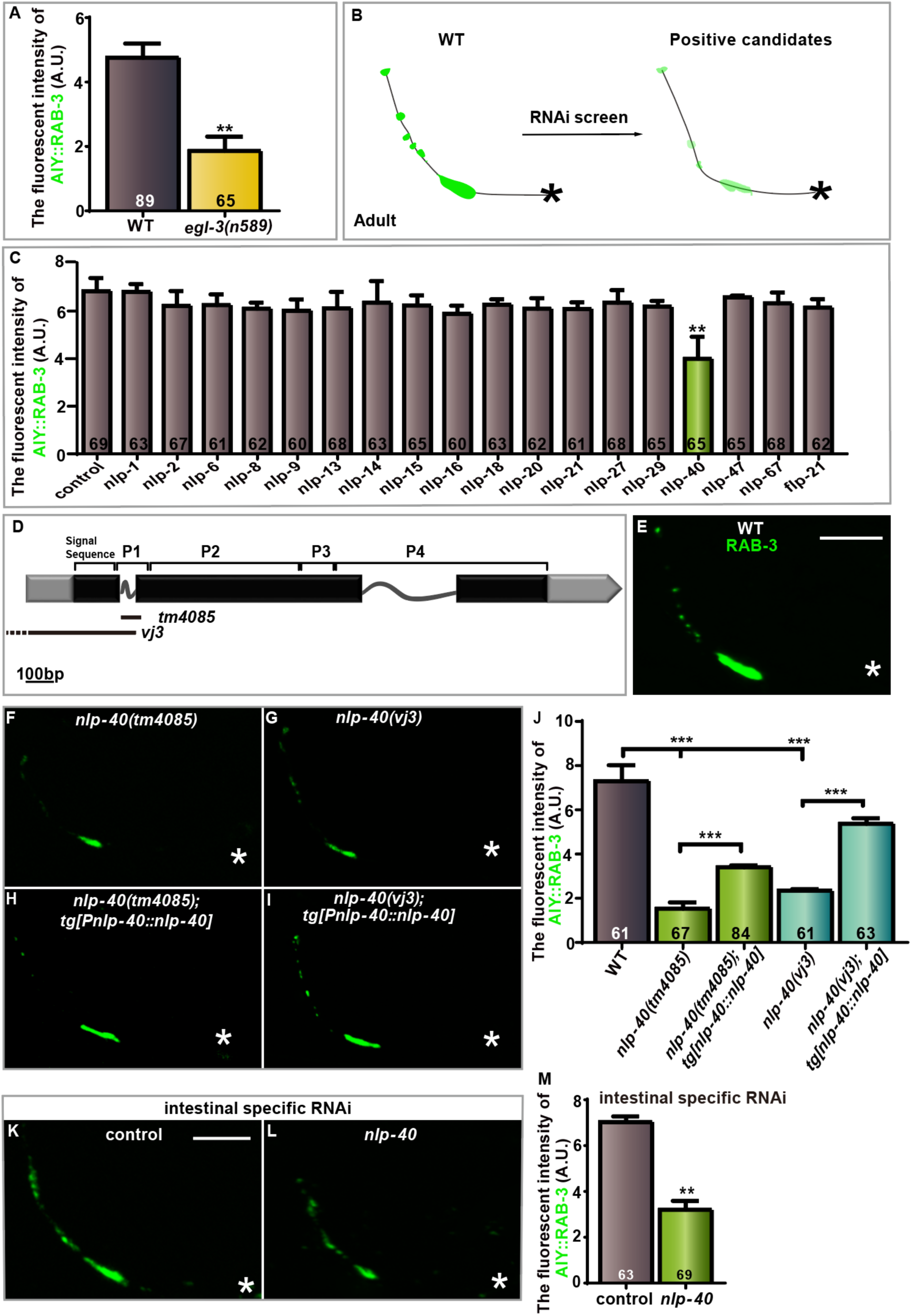
AIY presynaptic assembly requires intestinal expressed *nlp-40*. **(A)** Quantification of GFP::RAB-3 fluorescent intensity in wild-type and *egl-3(n589)* mutants. **(B)** A Schematic diagram describes the RNAi screen strategy to identify intestinal neuropeptides required for AIY synaptogenesis. **(C)** Quantification of GFP::RAB-3 fluorescent intensity for the indicated RNAi treatment. Candidate genes are intestinal expressed *nlp* or *flp*. **(D)** Diagram of *nlp-40* genomic structure. P1 to P4 indicates the predicted peptides encoded by *nlp-40*. The boxes and lines represent exons and introns. Black and gray indicate coding sequence and UTRs. The lines beneath indicate the deletion region. **(E-I)** Confocal images of AIY labeled with GFP::RAB-3 in wild-type (E), *nlp-40(tm4085)* (F), *nlp-40(vj3)* (G), *nlp-40(tm4085)* (H) or *nlp-40(vj3)* (I) with a wild type *nlp-40* transgene. The scale bar in (E) is 10μm and applies for (F-I). **(J)** Quantification of GFP::RAB-3 fluorescent intensity for the indicated genotypes. Transgenic data are averaged from at least two independent lines. **(K-L)** Confocal images of AIY synaptic marker GFP::RAB-3 for the indicated intestinal-specific RNAi treatment. The scale bar in (K) is 10μm and applies to (L). **(M)** Quantification for (K) and (L). For (A, C, J and M), data for each genotype are averaged from at least three biological replicates. The total number (N) of independent animals assayed is indicated in the bars. Statistical analyses are based on one-way ANOVA followed by Dunnett’s test (C and J) or two-tailed Student’s t-test (A and M). **p<0.01, ***p<0.001. Error bars represent SEM.

To confirm if AIY synaptogenesis requires *nlp-40*, we first examined the AIY synaptic phenotype in two independently isolated *nlp-40* mutant alleles, *tm4085* and *vj3* (Wang et al., 2013). *nlp-40(tm4085)* harbors an in-frame deletion that deletes the neuropeptide P1 and part of P2, an A70 to V mutation in the P3 and an early stop deleting the last three amino acid at the C-terminus of P4 (Fig. 2D and Supplementary Fig. 3). The *vj3* deletes the *nlp-40* promoter, the first exon and first intron (Fig. 2D)(Wang et al., 2013). Similar to the RNAi results, we observed a robust reduction of the AIY synaptic vesicle marker GFP::RAB-3 in both *nlp-40* mutant alleles (the GFP intensity was reduced by 79.01% and 67.49% respectively. P<0.001 compared to WT for both alleles; Fig. 2E-G, J). Similar results occurred with the active zone marker GFP::SYD-1 (78.97% reduction from the wild-type level. P<0.0001; Supplementary Fig. 4A-D, A’-C’). The AIY presynaptic defects in *nlp-40(tm4085)* and *nlp-40(vj3)* mutants were rescued by the wild type *nlp-40* transgene (the transgene increases the synaptic GFP intensity by 34.56% in *nlp-40(tm4085)* and by 50.15% in *nlp-40(vj3)*. P < 0.001 for both sets comparisons; Fig. 2H-J). Interestingly, overexpressing *nlp-40* in wild-type animals does not affect the AIY GFP::RAB-3 fluorescent intensity, suggesting that the role of *nlp-40* in AIY presynaptic assembly is not simply dosage dependent (Supplementary Fig. 5A-C). Based on these data, we conclude that that *nlp-40* is required for AIY synaptogenesis.

Next, we wanted to determine the NLP-40 action site. First, we examined the *nlp-40* expression pattern with both transcription and translational reporters and found *nlp-40* was high expressed in the intestine beginning at early embryonic stages, and the protein was observed both in the intestine and the coelomocytes, which is consistent with previous report (Supplementary Fig. 6A-E) (Wang et al., 2013). Then, using intestinal specific RNAi to knock down *nlp-40*, we observed a robust reduction of AIY presynaptic marker GFP::RAB-3, phenocopying the *nlp-40* mutants (the GFP::RAB-3 was reduced by 54.48%. P < 0.01; Fig. 2K-M). These data collectively suggest that *nlp-40* is expressed in the intestine and secreted to pseudocoelom. We posit intestinal NLP-40 likely travels to the nervous system and promotes AIY presynaptic assembly.

To address when *nlp-40* is required for the AIY presynaptic formation, we quantified the fluorescent intensity of AIY GFP::RAB-3 in *nlp-40(tm4085)* mutants at the larval L1, L4 and adult day 1 stage. We found the GFP intensity was dramatically reduced in the *nlp-40(tm4085)* mutants as of the newly hatched L1 stage throughout the adult stage (Supplementary Fig. 6F-H, 6F’-H’, 6I). These data suggest that intestinal *nlp-40* is required for AIY presynaptic assembly during embryonic development.

### *nlp-40* acts downstream of the Wnt signaling pathway to promote AIY presynaptic assembly

Thus far, we demonstrated that AIY presynaptic assembly requires both Wnt signaling and *nlp-40* from the intestine. To determine the nature of their interaction, we examined the genetic interaction between the genes encoding Wnt signaling components and *nlp-40* by building three sets of double mutants: *cwn-2(ok895);nlp-40(tm4085), cfz-2(ok1201);nlp-40(tm4085)* and *pop-1(hu9);nlp-40(tm4085)*. We found that the fluorescent intensity of the AIY presynaptic marker GFP::RAB-3 in all double mutants was similar to that in *nlp-40(tm4085)* single mutants, but slightly weaker than in Wnts single mutants (Fig. 3A). The epistatic effects of *nlp-40* to Wnt genes suggest that *nlp-40* acts downstream of the Wnt signaling pathway to regulate AIY presynaptic assembly.

**Fig. 3.**
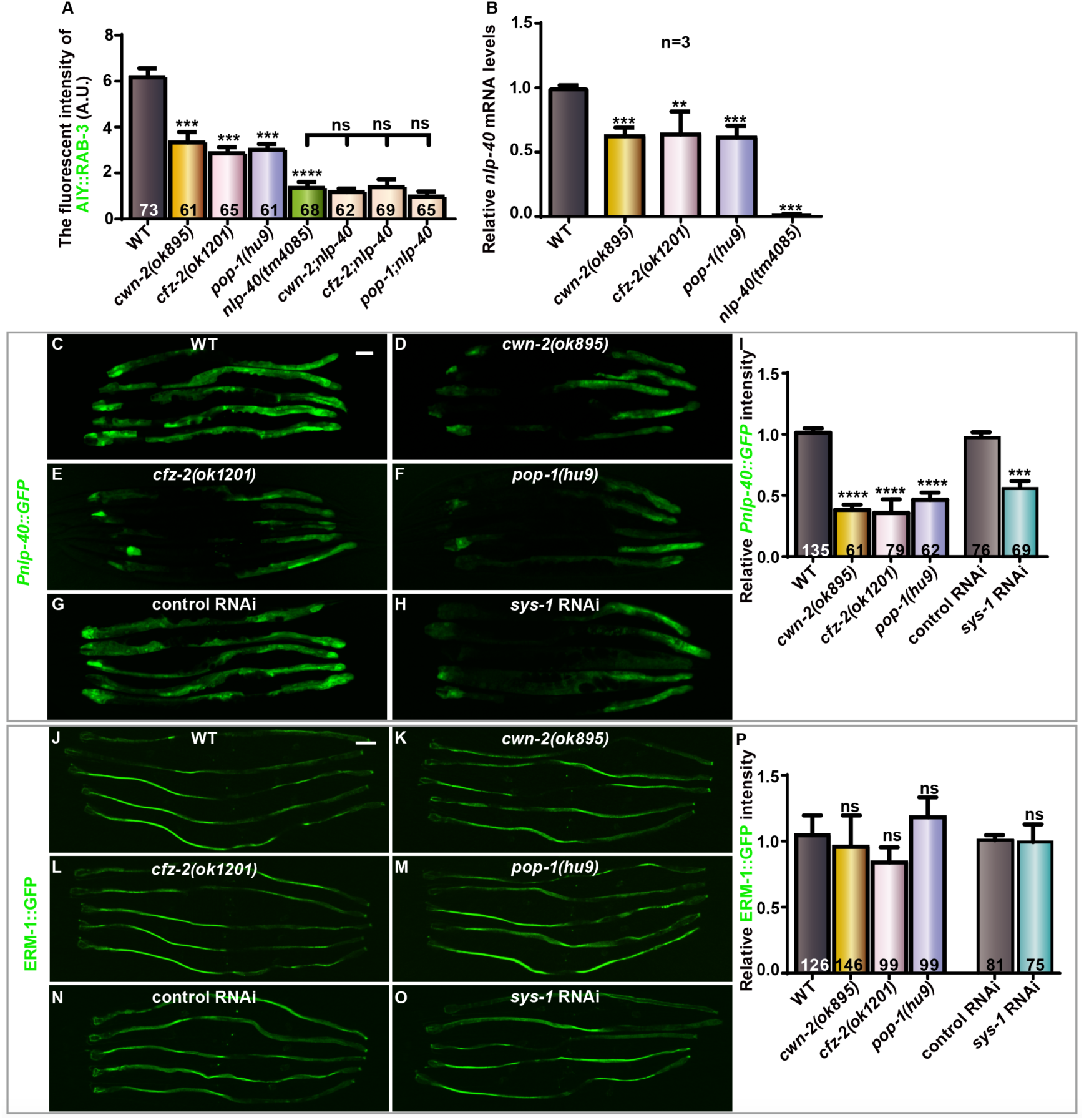
Wnt signaling regulates AIY presynaptic assembly by promoting *nlp-40* expression. **(A-B)** Quantification of the fluorescent intensity of AIY presynaptic marker GFP::RAB-3 and *nlp-40* mRNA level for the indicated genotypes. Data are averaged from three biological replicates. Statistical analyses are based on one-way ANOVA followed by Dunnett’s test. **p<0.01; ***p<0.001 as compared with wild-type. Error bars represent SEM. **(C-H)** Confocal images of the transcriptional *nlp-40* reporter (*Pnlp-40::GFP*) in wild-type (C), *cwn-2(ok895)* (D), *cfz-2(ok1201)* (E), *pop-1(hu9)* (F), and control or *sys-1* RNAi (G and H) animals at adult Day 1 stage. Scale bars: 50μm. **(I)** Quantification of P*nlp-40::GFP* fluorescent intensity for the indicated genotypes. **(J-O)** Confocal images of an intestinal-specific reporter ERM-1::GFP in wild-type (J), *cwn-2(ok895)* (K), *cfz-2(ok1201)* (L), *pop-1(hu9)* (M), and control or *sys-1* RNAi (N and O) animals at adult Day1 stage. The scale bar in (C) is 50μm and applies to the rest images. **(P)** Quantification of ERM-1::GFP fluorescent intensity for the indicated genotype. For (A, B, I and P) data are averaged from at least three biological replicates. The total number (N) of independent animals assayed is indicated in the bars. Statistical analyses are based on one-way ANOVA followed by Dunnett’s test and two-tailed Student’s t-test. ***p<0.001, ****p<0.0001, ns: not significant as compared to WT or control RNAi (p>0.05). Error bars represent SEM.

The above genetic data prompted us to test whether the Wnt signaling pathway regulates *nlp-40* expression. First, we quantified the *nlp-40* mRNA levels by quantitative reverse transcription PCR, and found that *nlp-40* mRNA levels were significantly reduced in *cwn-2(ok895), cfz-2(ok1201),* or *pop-1(hu9)* mutants compared to wild-type animals (the mRNA levels were reduced by 37.55%, 36.16% and 38.64% in *cwn-2(ok895), cfz-2(ok1201),* or *pop-1(hu9)* mutants. P<0.001, 0.01 and 0.001 respectively; Fig. 3B). Then, we quantified the *nlp-40* transcriptional reporter P*nlp-40::GFP* and observed a significant reduction of the GFP intensity in *cwn-2(ok895), cfz-2(ok1201)* or *pop-1(hu9)* mutants or *sys-1* RNAi animals (the GFP intensity was reduced by 62.38%, 64.65%, 53.98% and 45.76% in *cwn-2(ok895), cfz-2(ok1201)* or *pop-1(hu9)* mutants or *sys-1* RNAi animals. P<0.0001, 0.0001, 0.0001 and 0.001 respectively; Fig. 3C-I). To exclude the possibility that Wnt signaling affects the overall expression of intestinal genes, we examined the expression of intestinal specific *erm-1* and *vha-6* reporters (Gobel et al., 2004; Oka et al., 2001). Both reporters were not affected by genetic mutations or RNAi knockdown of the same set of Wnt signaling components (Fig. 3J-P and Supplementary Fig. 7A-F). Together, our data demonstrate that the Wnt signaling pathway promotes AIY presynaptic assembly by upregulating *nlp-40* in the intestine.

### Mature peptide P3 derived from NLP-40 is required for the AIY presynaptic assembly

*nlp-40* encodes a neuropeptide precursor protein, which contains a N-terminal signal peptide sequence and four predicted mature peptides (P1 to P4 in Fig. 4A)(Wang et al., 2013). To test which of the four peptides is essential for the AIY presynaptic assembly, we constructed four NLP-40 mutant variants (M1 to M4), each containing three mutant sites inactivating one corresponding mature peptide (Fig. 4A)(Wang et al., 2013). While the M1, M2 and M4 partially or fully rescued the AIY synaptic defect in *nlp-40(tm4085)* mutants, the M3 was not able to rescue at all (Fig. 4B-H, 4I). Those data indicate that the P3 is required for the AIY presynaptic assembly.

**Fig. 4.**
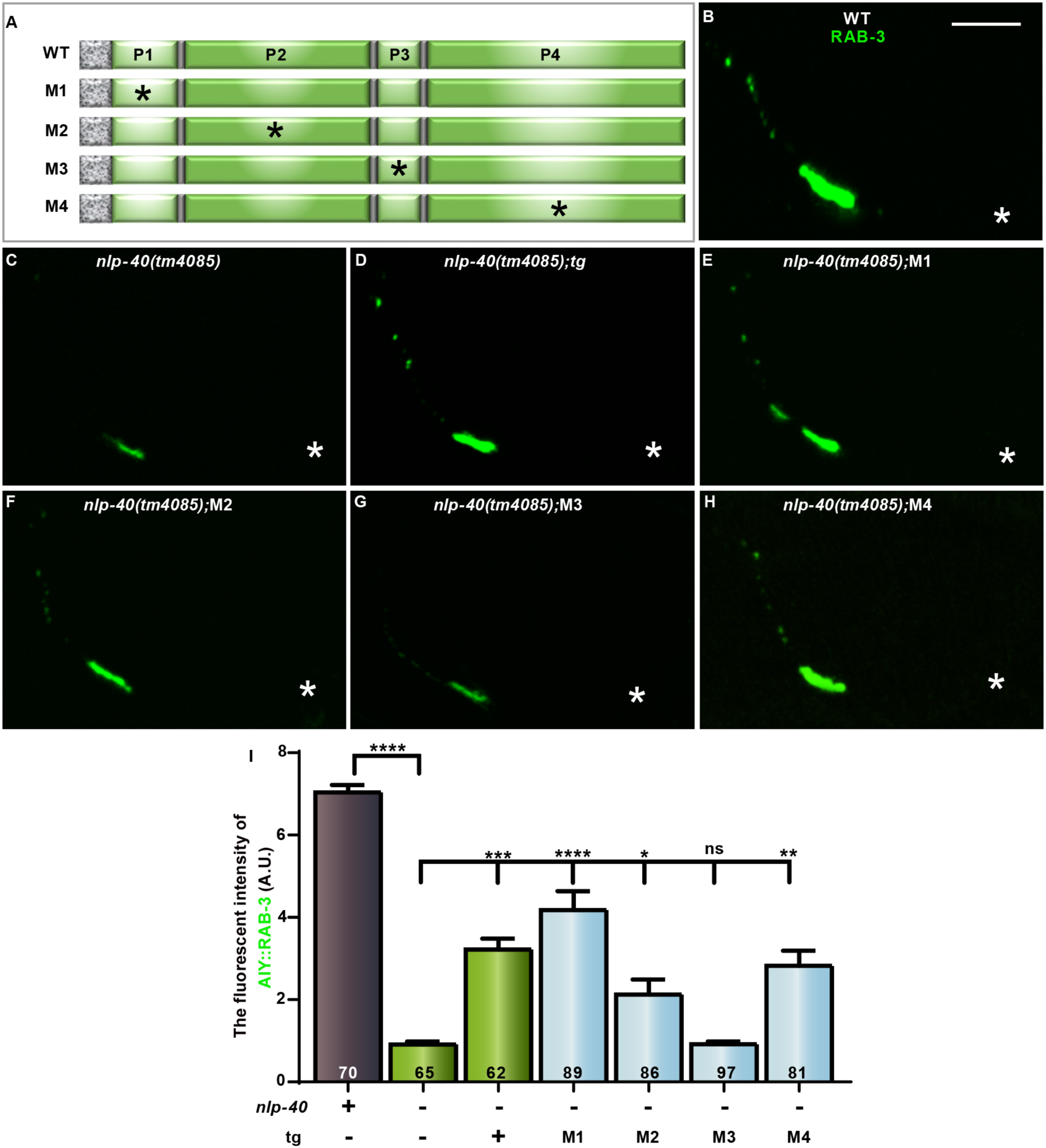
AIY presynaptic assembly requires peptide P3 derived from NLP-40. **(A)** Schematic of the neuropeptide precursor protein NLP-40. P1-P4 are predicted mature peptides derived from NLP-40 (Wang et al 2013), WT: *Pnlp-40::nlp-40*. Marble regions represent signal sequences, M1-M4 are missense mutations inactivating P1-P4 respectively. (M1 [ (24’ GLE) to AAA]; M2 [(62’ FVP) to AAA]; M3[(71’ WQP) to AAA]; M4[(109’PEE) to AAA]). **(B-H)** Confocal images of AIY GFP::RAB-3 in wild-type (B), *nlp-40 (tm4085)* (C), and the *nlp-40(tm4085)* with *Pnlp-40::nlp-40* (WT) (D), *Pnlp-40::nlp-40* (M1) (E), *Pnlp-40::nlp-40* (M2) (F), *Pnlp-40::nlp-40* (M3) (G), and *Pnlp-40::nlp-40* (M4) (H). The scale bar in (B) is 10μm and applies for (C-H). **(I)** Quantification of AIY GFP::RAB-3 intensity for the indicated genotypes. tg: P*nlp-40::nlp-40* cDNA transgene, +: wild type, -: mutant or without tg. Data are averaged from at least three biological replicates. M1-M4 indicate the mutant forms of NLP-40 as indicated in (A). Transgenic data are averaged from at least two independent lines. The total number (N) of independent animals assayed is indicated in the bars. Statistical analyses are based on one-way ANOVA followed by Dunnett’s test. ns: not significance (p>0.05), *p<0.05, **p<0.01, ***p<0.001, ****p<0.0001. Error bars represent SEM.

### The NLP-40 GPCR receptor AEX-2 acts cell-autonomously in the AIY to regulate presynaptic assembly

Recent studies found that AEX-2 serves as the NLP-40 receptor to regulate rhythmic behavior or anoxic survival (Doshi et al., 2019; Wang et al., 2013). To test whether *aex-2* is required for AIY presynaptic assembly, we examined the fluorescent intensity of AIY GFP::RAB-3 in *aex-2(sa3)* mutants (Fig. 5A), and found a robust reduction (76.84% reduction, P<0.0001; Fig. 5C, I). The reduction of the AIY presynaptic intensity in *aex-2(sa3)* mutants was rescued by a wild-type *aex-2* transgene (Fig. 5F, J). The data suggest that *aex-2* is required for AIY presynaptic formation.

**Fig. 5.**
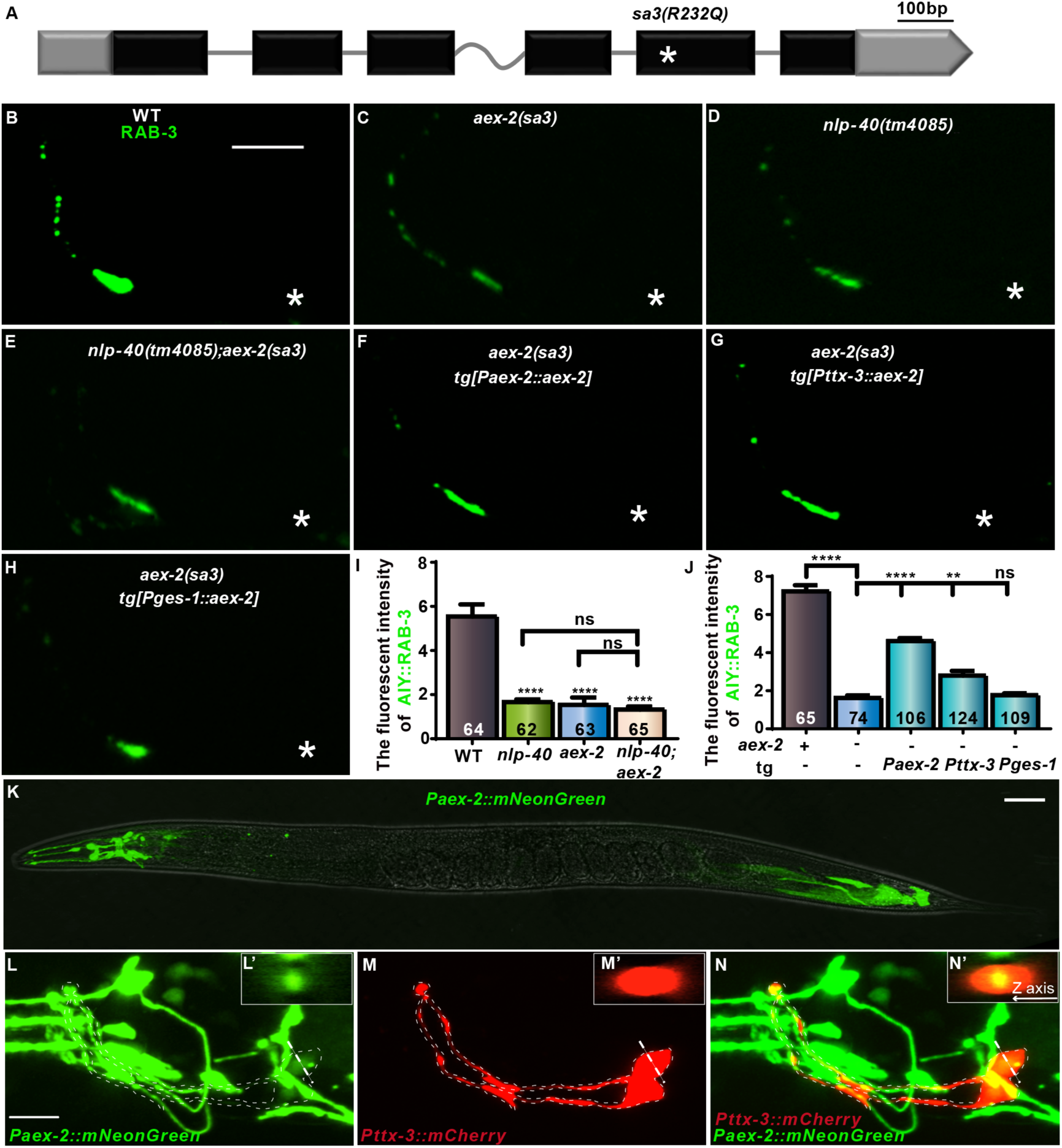
NLP-40 receptor AEX-2 is required for AIY presynaptic assembly. **(A)** Diagram of *aex-2* genomic structure. The boxes and lines represent exons and introns. Black and gray indicate coding sequence and UTRs. The red asterisk marks the location of the R>Q change in the *sa3* allele. **(B-H)** Confocal images of AIY GFP::RAB-3 in wild type (B), *aex-2(sa3)* (C), *nlp-40(tm4085)* (D) and *nlp-40(tm4085);aex-2(sa3)*(E) mutants and *aex-2(sa3)* with a wild-type *aex-2* transgene driven by the endogenous promoter(*Paex-2*) (F), AIY-specific promoter (*Pttx-3*) (G) or intestinal-specific promoter (*Pges-1*) (H). The scale bar in (B) is 10μm and applies to (C-H). **(I-J)** Quantification of AIY GFP::RAB-3 fluorescent intensity for the indicated genotypes. Data are averaged from at least three biological replicates. Transgenic data are averaged from at least two independent lines. The total number (N) of independent animals assayed is indicated in the bars. Statistical analyses are based on one-way ANOVA followed by Dunnett’s test. tg: *Paex-2::aex-2, Pttx-3::aex-2* or *Pges-1::aex-2* transgene, +: wild type; -: mutant or without tg, ns: not significance (p>0.05), **p<0.01, ****p<0.0001. Error bars represent SEM. **(K-N’)** Confocal images of *Paex-2*::mNeonGreen of whole animal with bright field (K), the head with *Paex-2*::mNeonGreen and *Pttx-3*::mCherry with green channel (L), red channel (M) and the merged channels (N). (L’-N’) are the cross sections corresponding to the dashed line sites. The scale bar in (K) is 50μm; the scale bar in (L) is 10μm and applies to (M and N).

To test the genetic interaction between *nlp-40* and *aex-2*, we quantified the AIY presynaptic marker GFP::RAB-3 in the *nlp-40(tm4085);aex-2(sa3)* double mutants. We found that the GFP intensity in the double mutants was similar to that in either single mutants (Fig. 5B-E, I), indicating that *nlp-40* and *aex-2* act in the same genetic signaling pathway to regulate AIY presynaptic assembly. The genetic interaction data is consistent with a model that AEX-2 acts as the NLP-40 receptor to regulate the AIY presynaptic assembly.

Next, we wanted to determine the *aex-2* action site. First, we built an *aex-2* transcription reporter, which showed that *aex-2* was mainly expressed in the nervous system with some in the posterior intestine (Fig. 5K), which is consistent with the previous report (Wang et al., 2013). When colabeled with the AIY cytoplasmic marker P*ttx-3::mCherry*, we noted that the *aex-2* reporter was co-labeled with the AIY marker (Fig. 5L-N’), indicating that *aex-2* is expressed in AIY interneurons. To address where *aex-2* acts, we expressed *aex-2* cDNA with AIY- or intestine-specific promoter in the *aex-2(sa3)* mutants, and found that only AIY-specific expression rescued the presynaptic defect (Fig. 5G, H, J). These data suggest that *aex-2* acts cell-autonomously in the AIY interneurons to regulate presynaptic formation.

### The Wnt-endocrine signaling regulates AIY activity

Neuropeptides can modulate ion channel and therefore neuronal activity (Davis and Stretton, 2001a; Matsushita and Arikawa, 1997; Rogers et al., 2001). NLP-40 has been shown to activate GABAergic neurons (Oliva and Inestrosa, 2015; Wang et al., 2013). To determine if Wnt/*nlp-40/aex-2* regulates AIY activity, we recorded the AIY calcium signaling using GCaMP6, a genetically encoded calcium indicator (Fig. 6A and 6A’)(Luo et al., 2014). In wildtype animals, the AIY automatically fires about six times (Wang et al., 2021). In *cfz-2(ok1201)*, *nlp-40(tm4085)* or *aex-2(sa3)* mutants, the fire frequency was reduced to half (Fig. 6B-C, Movie S1). However, the firing amplitude was not affected (Fig. 6D). These data suggest that Wnt-NLP-40 signaling pathway promotes AIY synaptic assembly probably through modulating neuronal activity.

**Fig. 6.**
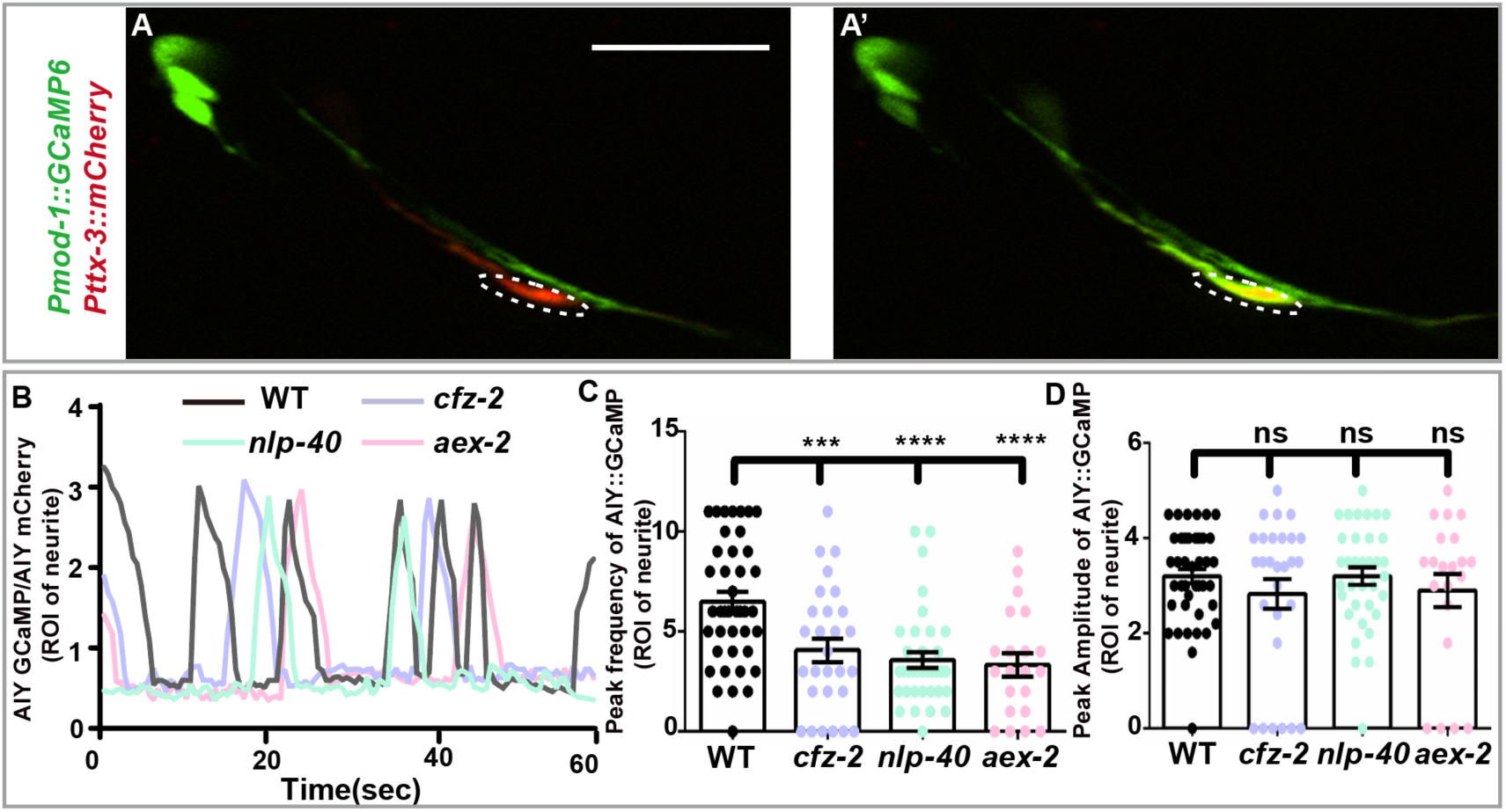
NLP-40 modulates AIY activity. **(A-A’)** Representative confocal images showing inactive (A) and active(A’) of AIY::GCaMP6 (with *mod-1* promoter) colabeled with mCherry (with *ttx-3* promoter) (red). Dashed ovals mark the AIY presynaptic enriched region where GCaMP was quantified. The scale bar in (A) is 10μm and applies to (A’). **(B)** Representative AIY calcium activity over 60 seconds for WT, *cfz-2(ok1201)*, *nlp-40(tm4085)* and *aex-2(sa3)* mutant animals. **(C-D)** Quantification of the peak frequency and amplitude of AIY::GCaMP6 in WT, *cfz-2(ok1201)*, *nlp-40(tm4085)* and *aex-2(sa3)* mutants. Each spot represents one animal. Data are averaged from at least three biological replicates. Statistical analyses are based on one-way ANOVA followed by Dunnett’s test. ns: not significance (p>0.05); ***p<0.001, ****p<0.0001. Error bars represent SEM.

### NLP-40 regulates synaptic protein trafficking

Neuropeptides and neurosteroids can regulate postsynaptic receptor trafficking (Lorenz-Guertin and Jacob, 2018). To test if NLP-40 regulates AIY presynaptic protein transports, we performed time-lapse to analyze the GFP::RAB-3 trafficking in the zone 1 region (Fig. 7A). We found that in the *nlp-40* mutants, the number of the GFP::RAB-3 puncta and the transport velocity of anterograde, but not retrograde, were significantly reduced (Fig. 7B-C”). Interestingly, the fluorescent intensity of GFP::RAB-3 in the soma was also significantly reduced, which is probably due to post transcriptional regulation since the promoter activity was not affected in *nlp-40* mutants (Fig. 7D and 7E). The decrease of GFP::RAB-3 could be due to the side effect of the inefficient trafficking. Collectively, those data suggest that NLP-40 regulates synaptic assembly most likely through modulating synaptic protein trafficking.

**Fig. 7.**
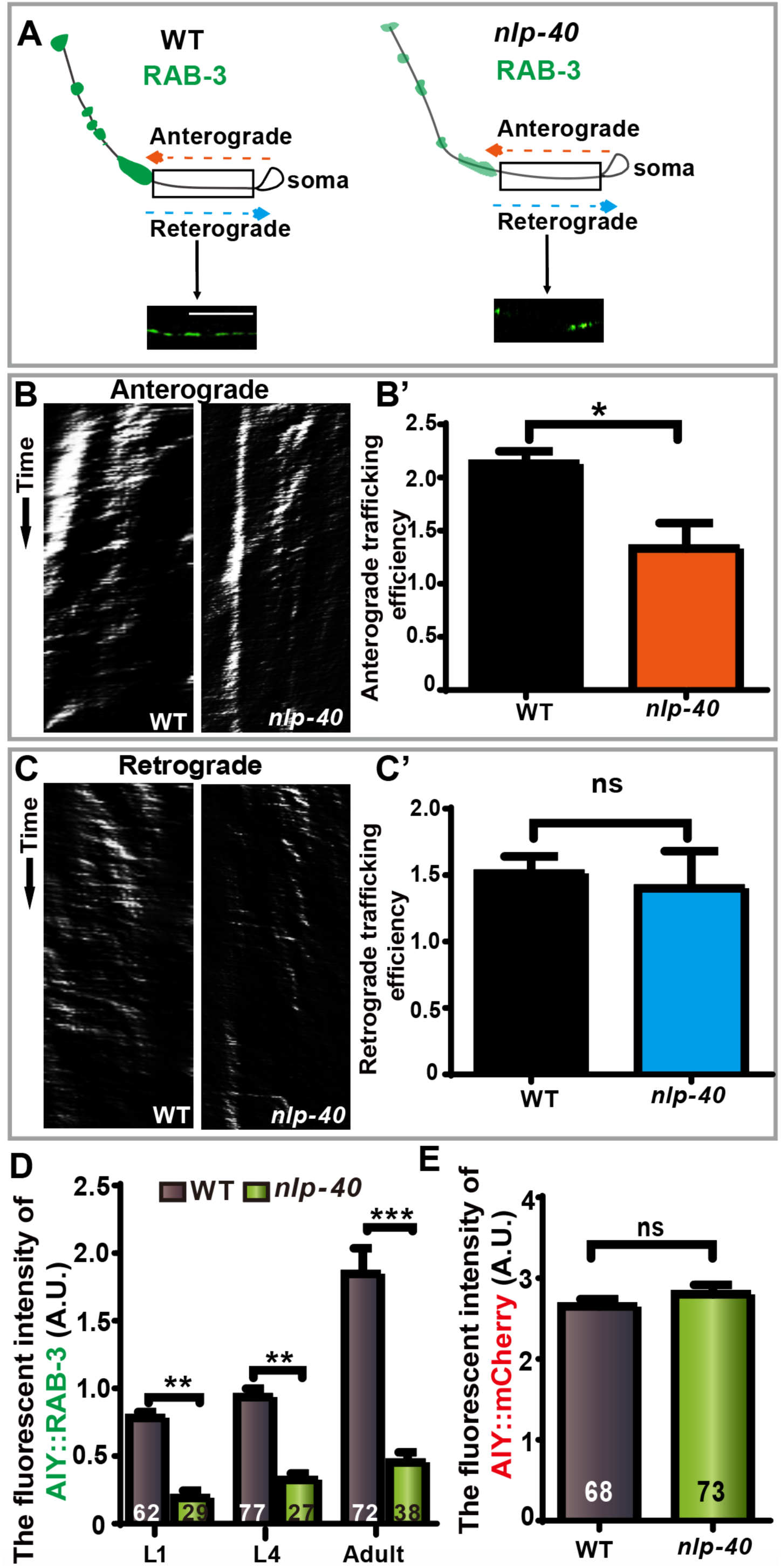
Neuropeptide/NLP-40 regulates synaptic protein anterograde trafficking. **(A)** A cartoon diagram of AIY GFP::RAB-3 in indicated genotypes. The AIY presynaptic sites are marked with green spots. Tracking zones are indicated with black boxes, and the representative confocal images are showed at the bottom. **(B, C)** The antero- and retro-grade AIY synaptic vesicle trafficking kymographs in WT and *nlp-40* mutants. **(B’, C’)** Quantification of the trafficking efficiency of GFP::RAB-3 corresponding to (B) and (C). Data are collected from at least three biological replicates. Each spot represents one individual. Statistical analyses are based on two-tailed Student’s t-test. ns: not significance (p>0.05), *p<0.05, Error bars represent SEM. **(D, E)** Quantification of the fluorescent intensity of P*ttx-3*::GFP::RAB-3 in the soma (D) (at L1, L4 and day1stage) and P*ttx-3::mCherry* (E) (at L3 stage) for the indicated genotypes. Data are collected from at least three biological replicates. The total number (N) of independent animals assayed is indicated in the bars. Statistical analyses are based on two-tailed Student’s t-test. ns: not significance (p>0.05), **p< 0.01, ***p<0.001. Error bars represent SEM.

### The Wnt/*nlp-40*/aex-2 signaling pathway is required for the presynaptic assembly in the nerve ring in general

Thus far, we have demonstrated that Wnt-NLP-40 signaling in the intestine is required for AIY presynaptic assembly. To determine if this regulation is general for presynaptic assembly or AIY specific, we first quantified GFP::RAB-3 in the nerve ring region in Wnt signaling mutants. Interestingly, the synaptic marker GFP::RAB-3 was significantly reduced in *cwn-2(ok895)*, *cfz-2(ok1201)* and *pop-1(hu9)* mutants (the intensity was reduced by 33.98%, 45.59% and 52.64% respectively, P<0.05, 0.01 and 0.001 as compared to the WT for mutants, respectively; Fig. 8A-E, H). Furthermore, we found that the reduction of the presynaptic marker in *cwn-2(ok895)* was rescued by a wild type transgene (Fig. 8I). Those data collectively suggest that the Wnt signaling is required for the presynaptic assembly in the nerve ring.

**Fig. 8.**
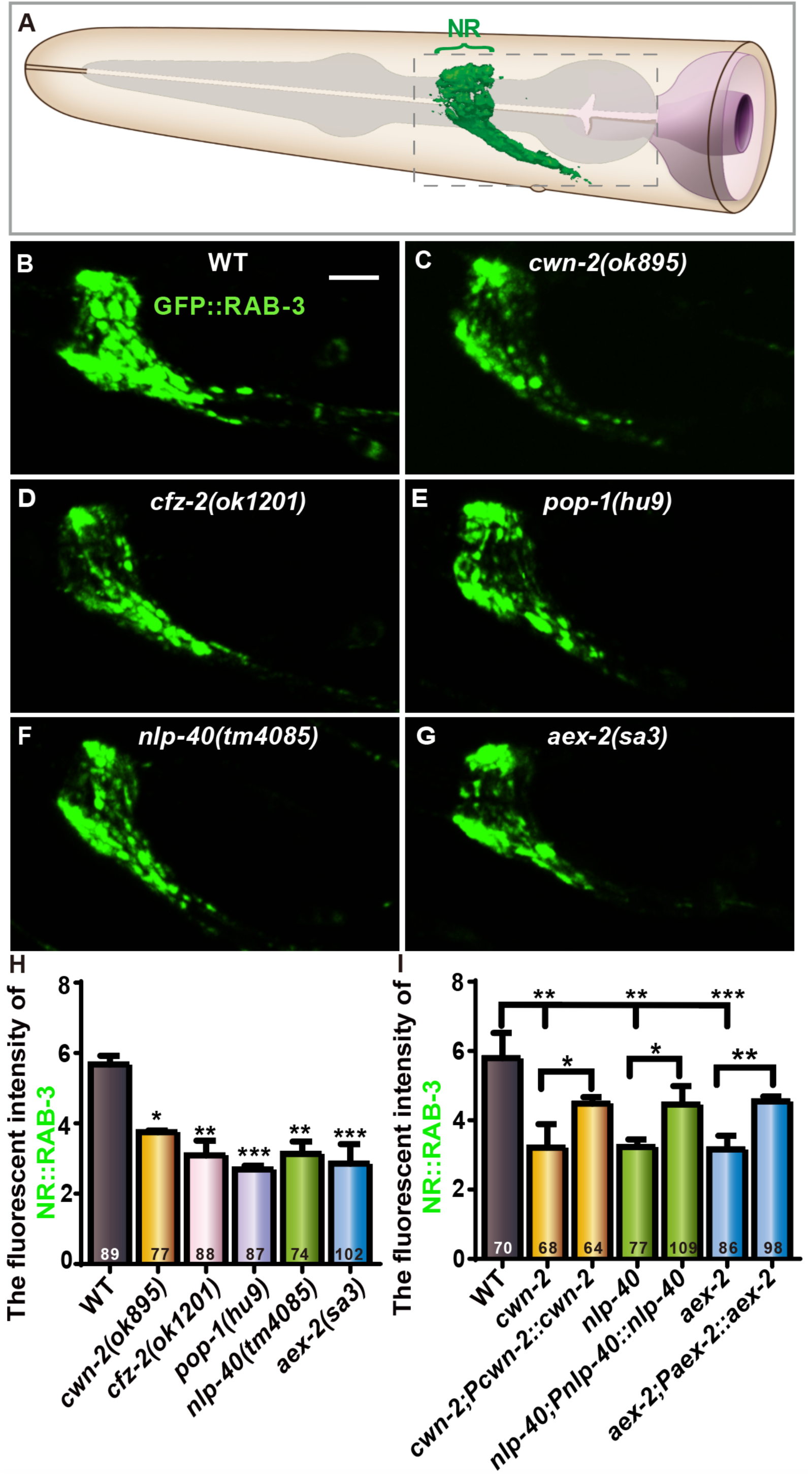
Synaptic assembly in the nerve ring requires Wnt-NLP-40 neuropeptide signaling. **(A)** A cartoon diagram of the *C. elegans* head. Green marks the presynaptic region in the nerve ring. **(B-G)** Confocal images of *Prab-3::GFP::rab-3* in the nerve region corresponding to the dashed box in (A)of wild type (B), *cwn-2(ok895)*(C)*, cfz-2(ok1201)*(D), *pop-1(hu9)*(E), *nlp-40(tm4085)*(F) and *aex-2(sa3)*(G) mutants. The scale bar in (B) is 10μm, applies to (C-G). **(H-I)** Quantification of the GFP intensity in the nerve ring for the indicated genotypes. Data are collected from at least three biological replicates. Transgenic data are averaged from at least two independent lines. The total number (N) of independent animals assayed is indicated in the bars. Statistical analyses are based on one-way ANOVA followed by Dunnett’s test and two-tailed Student’s t-test. *p<0.05, **p<0.01, ***p<0.001. Error bars represent SEM.

Next, we examined the presynaptic marker in *nlp-40* and *aex-2* mutants. We found that the GFP::RAB-3 intensity is dramatically reduced in *nlp-40(tm4085)* or *aex-2(sa3)* mutants (the intensity is reduced by 44.89% and 49.65% respectively, P<0.01 and 0.001 compared to WT; Fig. 8F, G and H). The reduction of the presynaptic marker was rescued by the corresponding wild type transgene (Fig. 8I). The results indicate that *nlp-40* and *aex-2* are required for presynaptic assembly in nerve ring.

## DISCUSSION

Overall, we demonstrate that a canonical Wnt signaling pathway that includes Wnt/CWN-2, Frizzled/CFZ-2, β-catenin/SYS-1 and TCF/POP-1 acts in the intestine to promote synaptic assembly in the *C. elegans* nerve ring, a structure analogous to mammalian brain. Wnt signaling exerts this effect by upregulating the expression of *nlp-40*, which encodes a conserved neuropeptide. NLP-40 promotes synaptic assembly through the neuronal expressed GPCR receptor, AEX-2 (Fig. 9). This gut Wnt-neuroendocrine signaling promotes synaptic assembly most likely through modulating neuronal activity. We therefore uncover a previously unidentified molecular mechanism in which the Wnt-neuroendocrine signaling pathway in the intestine regulates synaptic assembly. These results highlight the intrinsic role of gut in the neurodevelopment, and reveal a critical effect of novel gut-brain communication.

**Fig. 9.**
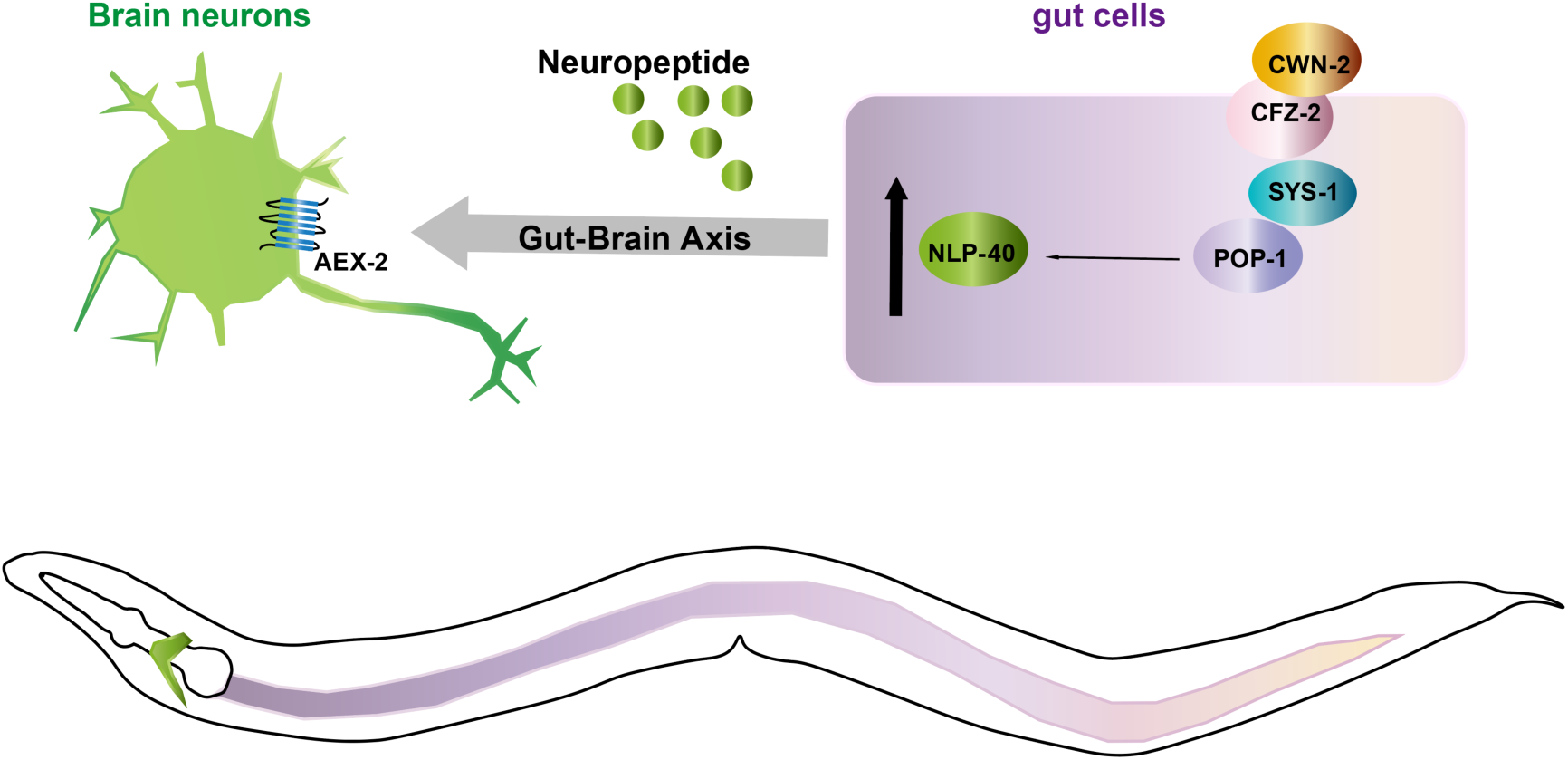
A model describing synaptic assembly mediated by Intrinsic Wnt-neuropeptide signaling from gut. A canonical Wnt signaling pathway including Wnt/CWN-2, Frizzled/CFZ-2, β-catenin/SYS-1 and TCF/POP-1 acts in the intestine to promote the synaptic assembly in the *C. elegans* nerve ring, a structure analogous to the mammalian brain. Wnt signaling promotes the expression of *nlp-40*, which encodes a conserved neuropeptide. NLP-40 secreted from intestine triggers the synaptic assemble through the neuronal expressed receptor GPCR/AEX-2.

### Wnt-Neuropeptide axis regulates synaptic assembly

The genes encoding Wnt signaling pathway components and their roles in synaptogenesis are highly conserved in *C. elegans* (He et al., 2018; Park and Shen, 2012). Wnt signaling plays critical roles in synaptic development. However, the underlying mechanisms are not completely understood. In this study, we demonstrated that intestinal Wnt signaling promotes synaptic assembly in the nerve ring. This Wnt signaling promotes synaptic assembly by upregulating the expression of the intestinal secreted neuropeptide, NLP-40, which acts through the neuronal expressed GPCR, AEX-2.

Wnt ligands act either in the nervous system or in non-neuronal tissues to regulate synaptic assembly or plasticity. However, the receptors or downstream components have only been shown to act in neurons or their target cells. For example, the Wnt ligands, LIN-44 and EGL-20, expressed in the tail suppress presynaptic assembly through the frizzled receptor LIN-17 or MIG-1 in DA8, DA9 and DB7 neurons (Klassen and Shen, 2007; Mizumoto and Shen, 2013). Presynaptic CWN-2 and LIN-44 regulate activity-dependent postsynaptic localization of the acetylcholine receptor ACR-16 mediated by the LIN-17 and CAM-1 receptors in the postsynaptic muscle cells (Jensen et al., 2012; Pandey et al., 2017). Through β-catenin and TCF/LEF transcription factors, the canonical Wnt signaling regulates the abundance of the postsynaptic glutamate receptor GLR-1(Dreier et al., 2005). Here, we found that a canonical Wnt signaling in the intestine promotes synaptic assembly in the *C. elegans* brain through the neuropeptide-like peptide NLP-40.

NLP-40 belongs to the neuropeptide-like proteins (NLPs) family and is required for rhythmic defecation and anoxia survival (Doshi et al., 2019; Husson et al., 2005; Wang et al., 2013). Neuropeptides are small bioactive signal peptides including insulin-like peptides (INS), FMRFamide-related peptides (FLPs) and neuropeptide-like proteins (NLPs) (Li and Kim, 2008). Those peptides are expressed both in the nervous system and non-neuronal tissues with a broad spectrum of biological functions (Chew et al., 2018; Hu et al., 2011; Hu et al., 2015; Li and Kim, 2008). For example, INSs regulate aversive learning behavior (Lee and Mylonakis, 2017) and feeding state-dependent gene expression in ADL neurons (Gruner et al., 2016). FLPs modulate the locomotion and body wave and head search behaviors (Chikka et al., 2016; Davis and Stretton, 2001b; Li, 2005; Reinitz et al., 2000). Similarly, NLPs regulate locomotion, body posture, rhythmic behaviors (Couillault et al., 2004).

The role of Wnt in synaptic development is highly conserved in vertebrates (Budnik and Salinas, 2011; He et al., 2018; Park and Shen, 2012). For example, Wnt7a and Wnt5a play critical roles in pre- or postsynaptic development in mammals (He et al., 2018). Although the role of Wnt-neuropeptide in synaptic development was previously not reported, The Wnt/β-catenin has been shown to regulate the expression of gut hormone/neuropeptides (Garcia-Jimenez, 2010; Garcia-Martinez et al., 2009; Kim et al., 2015), suggesting that Wnt-neuropeptide interactions are evolutionally conserved. In addition, hormones/neuropeptides are also involved in synaptic assembly and function in mammals (Corvino et al., 2015; D’Ercole et al., 2002; Garcia et al., 2014; Wang et al., 2015). Collectively, these studies suggest that the function of Wnt-neuropeptide axis in synaptic assembly is most likely conserved in vertebrates.

### An intrinsic role of gut in synaptic development

Using tissue-specific RNAi, tissue-specific rescue and transgenic GFP reporter experiments, we demonstrated that the Wnt-NLP-40 axis acts in the gut to regulate presynaptic assembly of nerve ring neurons. Further, we showed that the effect of the Wnt-NLP-40 axis on synaptic assembly at the newly hatched L1 stage, suggesting it acts during embryogenesis before feeding. We propose that the Wnt-NLP-40 axis acts through a stimulus-independent, intrinsic developmental program.

The gut-brain communication plays an important role in coping with unfavorite environmental conditions. For example, under stress conditions, intestinal SKN-1 regulates locomotion by modulating synaptic transmission at the neuromuscular junctions (D’Ercole et al., 2002; Staab et al., 2013). Insulin-like peptides INSs expressed in the intestine can regulate pathogen avoidance or feeding behavior through modulating the function of nervous system (Gruner et al., 2016; Lee and Mylonakis, 2017). Intestinal infections can affect the host’s learning and behavior by neuroendocrine signaling pathways (Singh and Aballay, 2019). Intestinal immune responses can also protect neurons from degeneration (Chikka et al., 2016). However, the intrinsic role of gut in the brain development is largely unexplored, and here we provide a molecular mechanism underlying this novel gut-brain interaction.

In vertebrates, the gut-brain-microbiota axis plays a vital role in mediating brain development and brain function, such as locomotion, exploratory and risk-taking behaviors and spatial memory (Bercik et al., 2011; Diaz Heijtz et al., 2011; Gareau et al., 2011; Neufeld et al., 2011). Endocrine is a highly conserved signaling pathway mediated the gut-brain communication (Farzi et al., 2018; Matafome et al., 2017; Sun et al., 2018). Although it is unknown if gut-brain axis plays any intrinsic role in neural circuit formation in vertebrates, we posit that the signal pathway regulating synaptic formation is probably conserved given the conservation of gut-brain axis, Wnt signaling, neuroendocrine signaling.

### Spontaneous neural activity regulates synaptic assembly

It is well known that neuroactivity regulates synaptic formation and plasticity (Hooks and Chen, 2020; Pan and Monje, 2020). The neuronal activity can regulate the trafficking of synaptic components (Lorenz-Guertin and Jacob, 2018; Sears and Broadie, 2017; Tiruchinapalli et al., 2003). Interestingly, here we found that Wnt-neuroendocrine signaling pathway is required for AIY spontaneously firing and synaptic vesicle trafficking, suggesting that the intrinsic gut signaling pathway most likely regulates synaptic formation through modulating neuronal activity.

Neuronal spontaneous activity is common during embryogenesis and plays critical roles in neuronal development including neurogenesis and circuit formation (Blankenship and Feller, 2010; Moody and Bosma, 2005). Based on our finding and studies from other groups, we speculate that the role of neuronal spontaneous activity in synaptogenesis is conserved.

Collectively, our findings not only provide mechanistic insights into the synaptic assembly, but also uncovered a novel gut-brain interaction.

## MATERIALS AND METHODS

### Strains and maintenance

All *C. elegans* strains were grown with *E. coli* OP50 on standard NGM plates at 21°C. A detailed information of all the strains is listed in Supplementary Table S1. Animals were used in this study at adult day 1 stage unless specified.

### Plasmids and transformation

Constructs were made either with pSM (derivation of pPD49.26) (Mello and Fire, 1995; Shen and Bargmann, 2003) or L4440 vector (derivation of pPD129.36) (Kamath and Ahringer, 2003). Detailed information is described in Table S2.

Four NLP-40 mutant variants (M1 to M4, corresponding to the defective P1-P4 respectively) were constructed with three consecutive residues replaced with alanine in one of the predicted mature peptides. (M1 [ (24’ GLE) to AAA, we change the (70’ ggtctcgag) to (gctgccgcg)]; M2 [(62’ FVP) to AAA, (184’ tttgttcca) to (gctgctgca)]; M3[(71’ WQP) to AAA, (211’ tggcagccg) to(gcggcggcg)]; M4[(109’PEE) to AAA, (325’ccggaggaa) to (gcggcggca)]) (Wang et al., 2013). Detailed information is described in Table S2.

Transgenic strains were generated by microinjections as previously described (Mello and Fire, 1995). We used P*hlh-17::mCherry*(20ng/μl), P*unc-122::gfp*(20ng/μl) or P*ttx-3::sl2::mCherry* (20ng/μl) as co-injection markers. Detailed information is described in Table S2.

### RT and qRT-PCR

qRT-PCR was performed as the previous study (Shen et al., 2012). In brief, for mRNA, total RNA was prepared by traditional Trizol extraction methods from ∼100 worms. cDNA was subsequently generated by GoScript^TM^ Reverse Transcription system for RT-qPCR (Bio-Rad). qRT-PCR was performed with 2xNovoStart®SYBR qPCR SuperMix Plus (Novoprotein) on a CFX384 Touch^TM^ Real-Time PCR Detection System (Bio-Rad). Three technical replicates were performed in each reaction. *nlp-40* was used as reference for mRNA quantification, the results were from at least three biological replicates. qRT-PCR primers for *nlp-40* were designed and was listed in Table S2.

### RNA interference

RNAi constructs were transformed into HT115. RNAi feeding experiments were performed based on the previous study (Fraser et al., 2000). For tissue-specific RNAi experiment, we used the worm strain FDU2851(*sid-1(qt9); alxIs9[Pvha-6::sid-1::sl2::GFP]; olaIs12[Pttx-3::mCherry::rab-3; Phlh-17::GFP])* and FDU3111(*sid-1(pk3321); uIs69[Punc-119::sid-1; Pmyo-2::mCherry]; wyIs45[Pttx-3::GFP::rab-3; Punc-122::RFP])* for intestine and neuronal specific RNAi respectively. To screen the intestinal neuropeptide encoding genes through RNAi knockdown for the AIY presynaptic regulators, we transferred ten synchronized day 1 adults to the RNAi plates, allow them to lay eggs for two hours, then remove those adults. Then, we quantified the AIY synaptic fluorescent intensity at the adult day 1 stage.

### Calcium imaging

The animals of FDU424(*olaIs17 [Pmod-1::GCaMP6; Pttx-3::mCherry; Punc-122::dsRed]*) and FDU3604 (*nlp-40(tm4085)*;*olaIs17 [Pmod-1::GCaMP6; Pttx-3::mCherry; Punc-122::dsRed]*) were used for AIY calcium imaging. The P*ttx-3::mCherry* were used as an internal control. Calcium imaging of adult day 1 animals was performed on 10% agarose pads immobilized with 0.1 µm polystyrene beads and covered with glass coverslip.

Images were captured with Andor Dragonfly Spinning Disc Confocal Microscope with 60X objectives, 488 nm (for GFP) or 561 nm (for mCherry) laser. Individual animals were imaged for 60s at a rate of 2 Hz. Data were analyzed using custom written scripts in R Studio, and the images rotation and brightness/contrast were processed with Adobe photoshop CC. The 60s video was exported with Imaris 4.0.

### Live image acquisition and Kymograph analysis

Animals at L3 stage were anesthetized on 3% agarose pads with 100 mM muscimol. All images were captured with Andor Dragonfly Spinning Disc Confocal Microscope with 60X objectives. The exposure time is 400ms and 800ms for wild-type and *nlp-40* mutants respectively. Time-lapse movies were acquired continually with 1s interval for 5min. For quantification, the time-lapse movies were processed by Fiji and KymographDirector2.1 (Fig. 7B, 7C). Firstly, the movies were imported into Fiji to get the Kymograph images with Kymograph clear 2.0. Each kymograph was processed with KymographDirector2.1 to generate the final data (GraphPad Software), and the intensity threshold was set based on the background.

### Fluorescence microscope and confocal imaging

For general microscopy experiments, worms were anesthetized on 3% agarose pads with 50 mM muscimol. Images were captured with Andor Dragonfly Spinning Disc Confocal Microscope with 10X or 40X objectives, 488 nm (for GFP) or 561 nm (for mCherry) laser. The fluorescence intensity was quantified with Imaris 4.0, and the images rotation and brightness/contrast were processed with Adobe photoshop CC.

### Quantification and statistical analysis

We quantified the fluorescence intensity using Imaris. The density obtained by cropping the whole AIY interneuron zone (showed in Fig. 1A dashed box) and the whole intestinal zone and NR zone (showed in Fig. 7A dashed box) in Imaris. To monitor the temporal role of indicated genes, we quantified the F1 animals’ phenotype when F0 day1 animals’ eggs laid and were synchronized as 12, 20, 30, 42, 66 hr later, respectively. All quantified data were collected from at least three biological replicates. For data with transgenic animals, at least two independent lines were used. The statistical P values were measured using GraphPad Prism 6.0 (GraphPad Software). Statistical analysis between two groups were based on Student’s t-test (two-tailed), among more than two groups were based on one-way ANOVA followed by Dunnett’s test, as indicated in the Fig. legends. All quantitative data were collected blindly.

## ACKNOWLEDGMENTS

We thank ZF Altun and DH Hall for help with schematic in Fig. 1A’ and 8A. We thank Derek Sieburth (University of Southern California), Shiqing Cai, Yidong Shen (Chinese Academy of Sciences), Huanhu Zhu (Shanghai Tech), Zhao Qin (Tongji University), Di Chen (Nanjing University) and *Caenorhabditis elegans* Genetic Center (funded by NIH (P40 OD010440) for providing strains or plasmids. We thank members in Shao laboratory for comments on the project. We thank Min Jang, Ying Shi, Mi Zhou from IOBS facility core at Fudan University for providing technical support on image acquisition. This study was supported by the National Natural Science Foundation of China (Grant No. 31872762). We thank Life Science Editors for editing assistance.

## ABBREVIATIONS

GI: gastrointestinal
EECs: enteroendocrine cells
CCK: cholecystokinin
PYY: peptide tyrosine-tyrosine
GLP-1: glucagon-like peptide-1
RNAi: RNA interference
PC2: proprotein convertase
GPCR: G Protein-Coupled Receptor
TCF: T-cell specific transcription factor
NLPs: neuropeptide-like proteins
ILPs: insulin-like peptides
FLPs: FMRFamide-related peptides

## COMPLIANCE WITH ETHICS GUIDELINES

The authors declare no competing interests. All institutional and national guidelines for the care and use of laboratory animals were followed.

## AUTHOR CONTRIBUTIONS

YS and ZS conceived, designed the project. YS and LQ performed all experiments. YS, LQ and ZS analyzed data and interpreted the results. YS and ZS wrote the manuscript.

---

**Supplementary Fig. 1.**
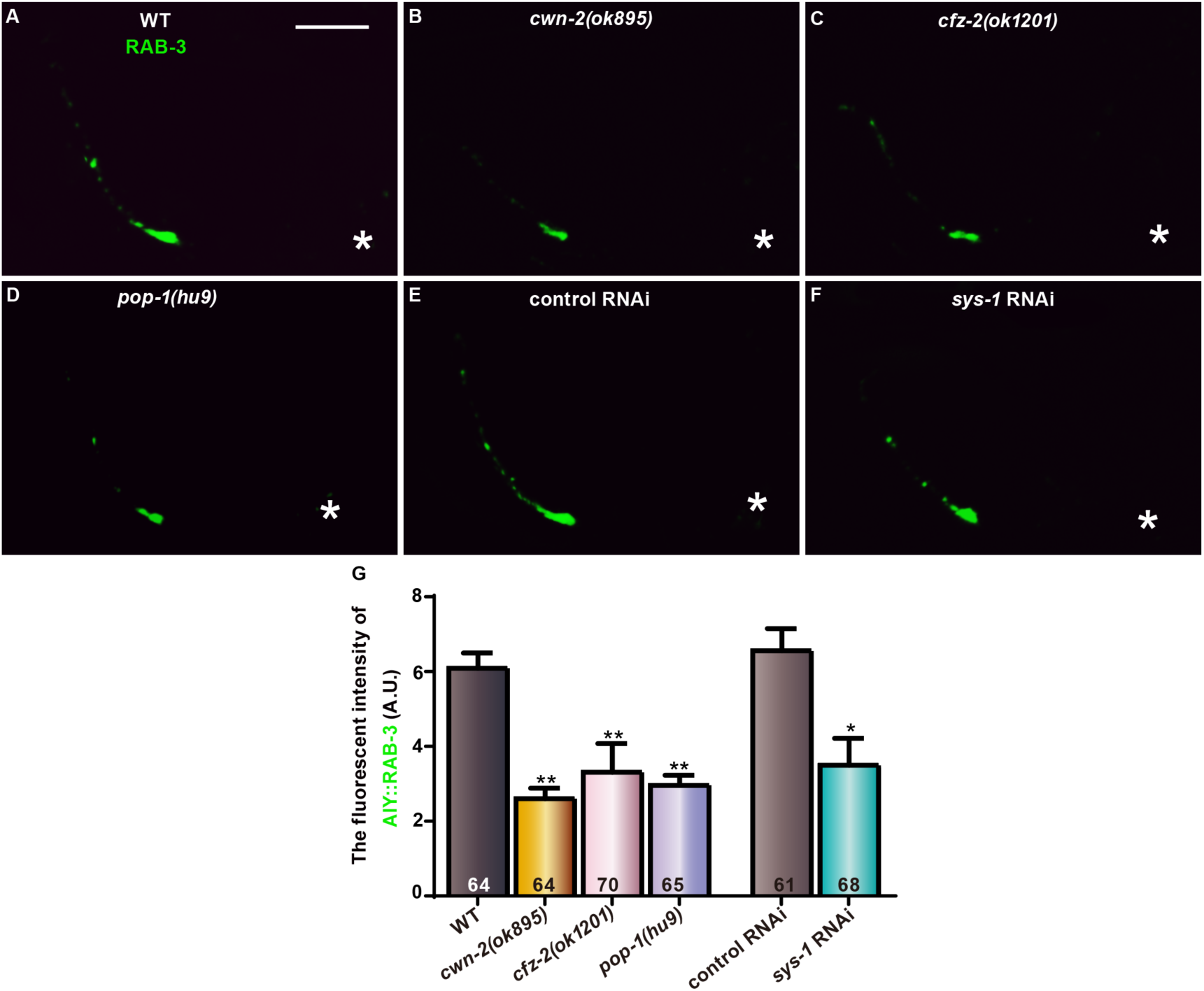
AIY presynaptic assembly requires canonical Wnt signaling components. **(A-F)** Confocal images of AIY synaptic marker mCherry::RAB-3 (pseudogreen) in wild type(A), *cwn-2(ok895)*(B), *cfz-2(ok1201)*(C), *pop-1(hu9)*(D) mutants, empty vector control (E) and *sys-1* RNAi (F) animals. The fluorescent intensity is significantly reduced in *cwn-2(ok895)*, *cfz-2(ok1201)*, *pop-1(hu9)*, or *sys-1* RNAi animals. Asterisks indicate AIY soma. The scale bar in (A) is 10μm, applying for all image panels. **(G)** Quantification of the fluorescent intensity of AIY RAB-3 for the indicated genotypes. Data are averaged from at least three biological replicates. The total number (N) of independent animals assayed is indicated in the bars. Statistical analyses are based on one-way ANOVA followed by Dunnett’s test (for mutants) or two-tailed Student’s t-test (for RNAi). *p<0.05, **p<0.01 as compared to WT or control RNAi. Error bars represent SEM.

**Supplementary Fig. 2.**
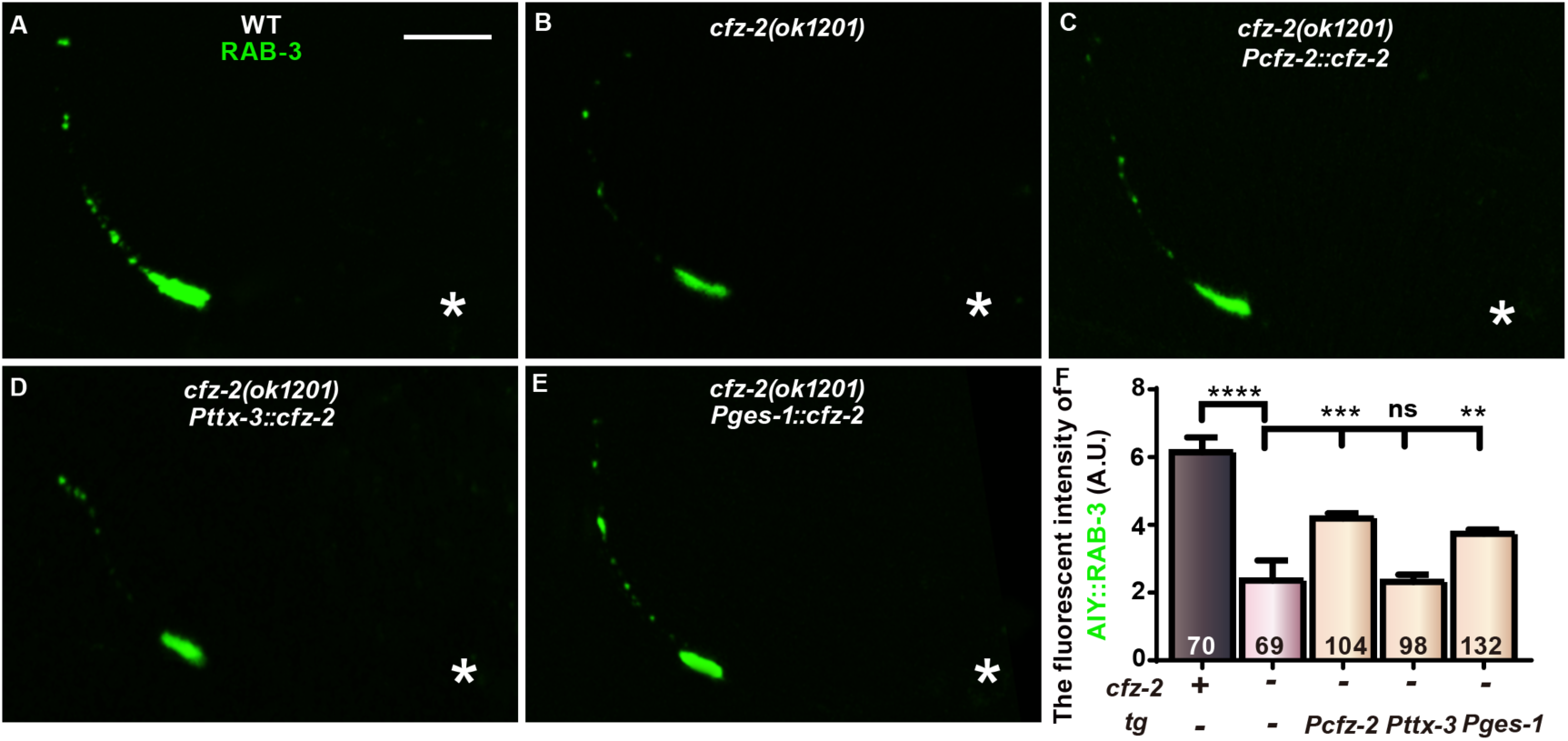
AIY presynaptic assembly requires intestinal *cfz-2*. **(A-E)** Confocal images of AIY GFP::RAB-3 in wild type (A), *cfz-2(ok1201)* (B), *cfz-2(ok1201)* with a wild-type *cfz-2* transgene driven by the endogenous promoter(*Pcfz-2*) (C), AIY-specific promoter (*Pttx-3*) (D) or intestine-specific promoter (*Pges-1*) (E). The scale bar in (A) is 10μm and applies to (B-E). **(F)** Quantification of AIY GFP::RAB-3 fluorescent intensity for the indicated genotypes. Data are averaged from at least three biological replicates. Transgenic data are averaged from at least two independent lines. The total number (N) of independent animals assayed is indicated in the bars. Statistical analyses are based on one-way ANOVA followed by Dunnett’s test. ns: not significant (p>0.05), **p<0.01, ***p<0.001, ****p<0.0001. Error bars represent SEM.

**Supplementary Fig. 3.**
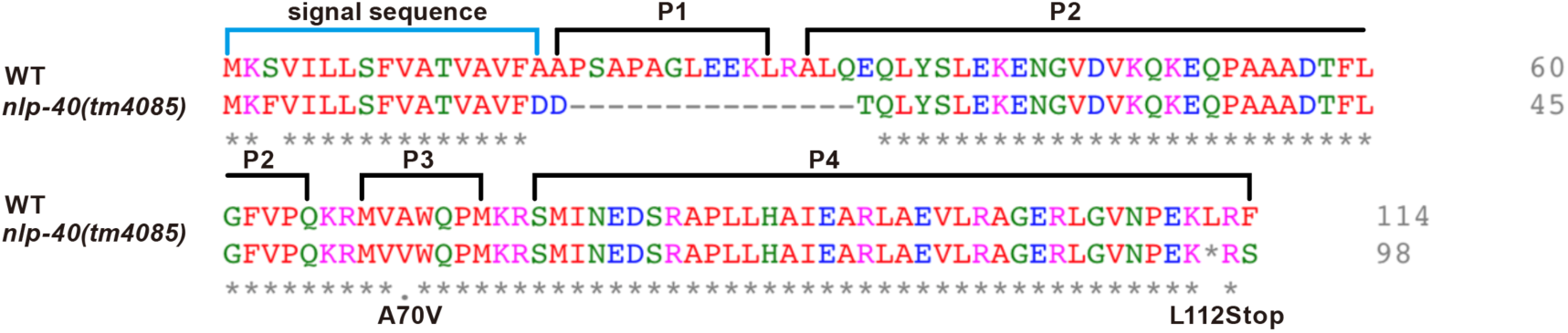
NLP-40 protein sequence alignment between wild-type and *nlp-40(tm4085)* The image represents the protein alignment between WT and *nlp-40(tm4085)*. P1 to P4 indicates the predicted mature products of NLP-40. Stars and dots indicate identity and similarity.

**Supplementary Fig. 4.**
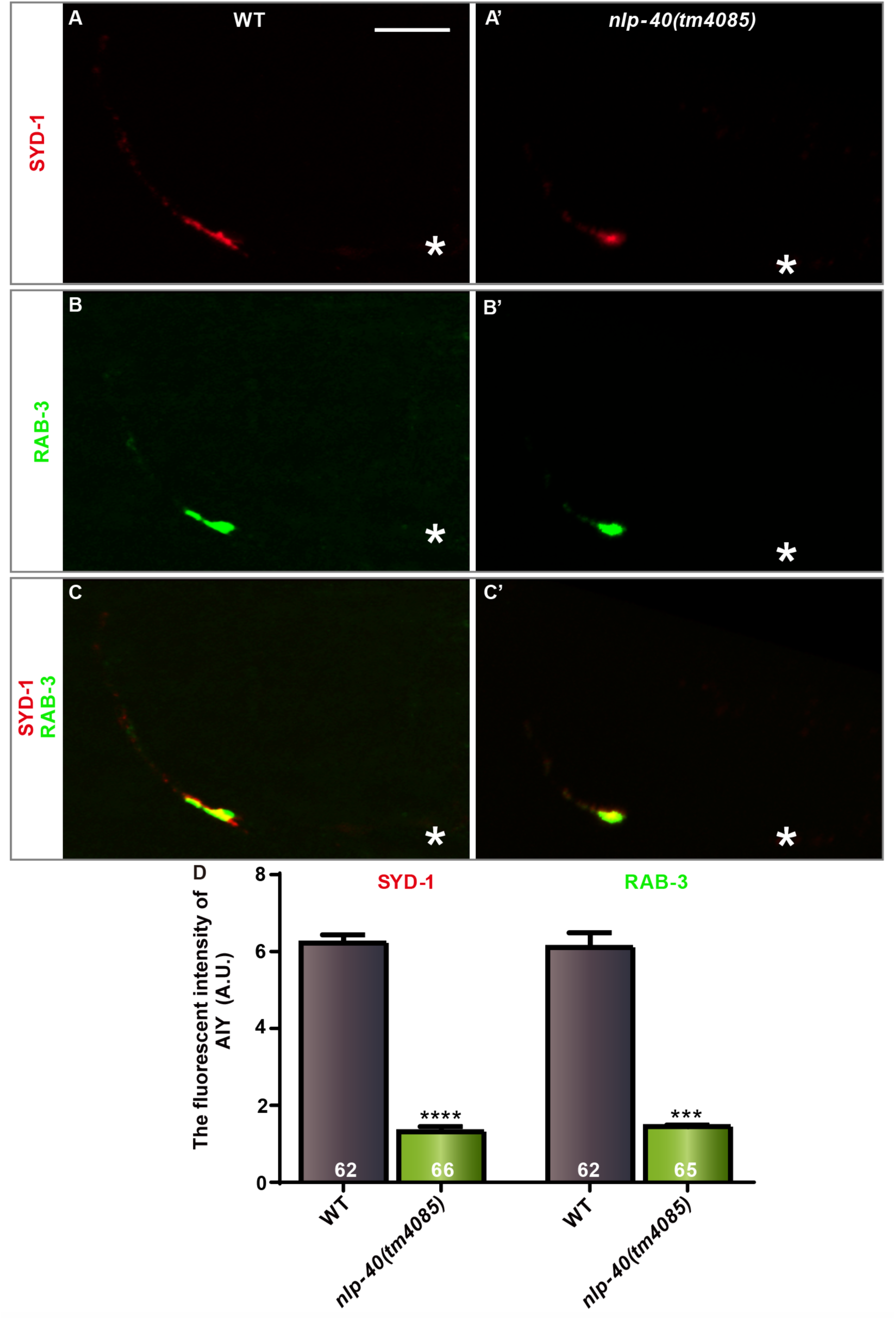
AIY presynaptic assembly requires *nlp-40*. **(A-C’)** Confocal micrographs of the AIY synaptic active zone GFP::SYD-1 (pseudo-red) and synaptic vesicle marker mCherry::RAB-3 (pseudo-green) in wild type (A-B) and *nlp-40 (tm4085)* mutants (A’-B’). Those two presynaptic markers are colocalized, as indicated in the merged images (C-C’). The scale bar in (A) is 10μm, applying to all image panels. **(D)** Quantification of the fluorescent intensity of AIY SYD-1 or RAB-3. Data are averaged from at least three biological replicates. The total number (N) of independent animals assayed is indicated in the bars. Statistical analyses are based on two-tailed Student’s t-test. ***p<0.001, ****p<0.0001 as compared to WT. Error bars represent SEM.

**Supplementary Fig. 5.**
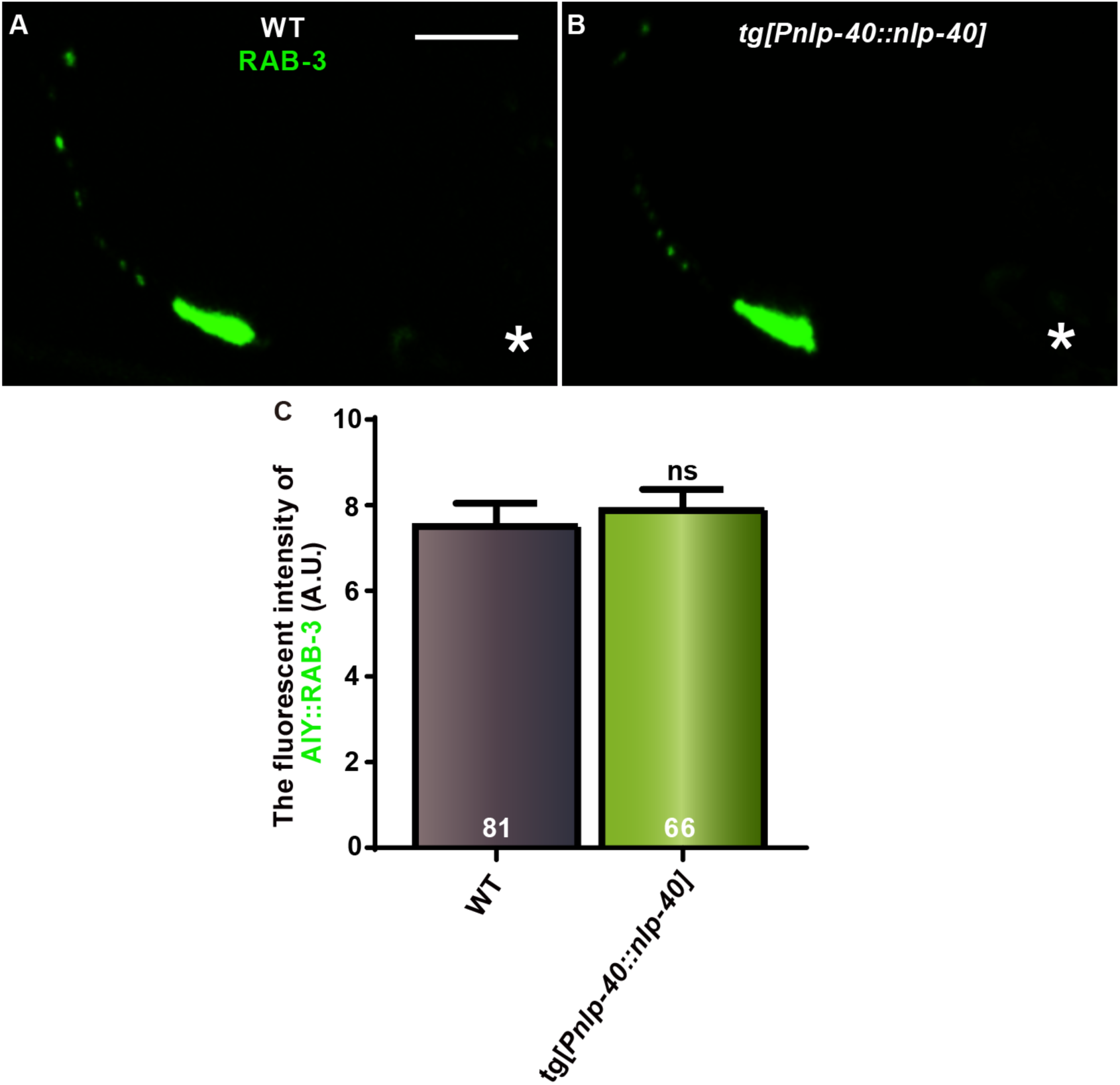
*nlp-40* overexpression has not effect on the AIY presynaptic assembly. **(A-B)** Confocal images of the AIY GFP::RAB-3 (A-B). The scale bar in (A) is 10μm, applying to (B). **(C)** Quantification of the fluorescent intensity of AIY RAB-3 for the indicated genotypes. Data are averaged from at least three biological replicates. The total number (N) of independent animals assayed is indicated in the bars. ns: not significant. Statistical analyses are based on two-tailed Student’s t-test. Error bars represent SEM.

**Supplementary Fig. 6.**
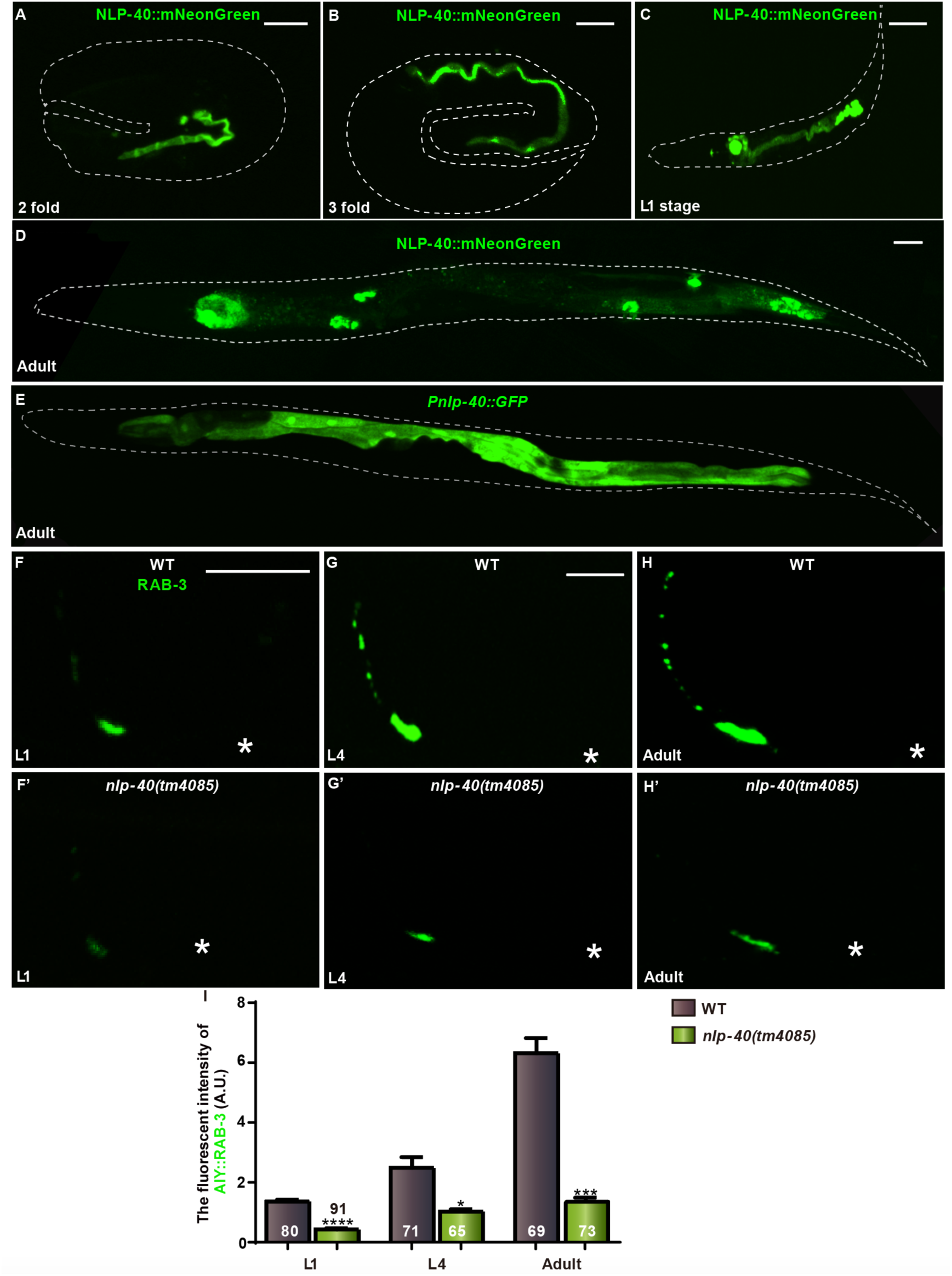
*nlp-40* is required for AIY synaptogenesis during embryonic and postembryonic development. **(A-E)** Confocal images of expression pattern and subcellular localization of embryonic stage, adult stage animals labeled with NLP-40::mNeonGreen (A-D) or GFP (E). Scale bars in (A-C) are 10μm, and the one in (D) is 50μm, applying to (E). **(F-H’)** Confocal micrographs of the AIY GFP::RAB-3 at L1 (F-F’), L4 (G-G’) and adult day 1 (H-H’) in wild type (F-H) and *nlp-40(tm4085)* mutant (F’-H’) animals. Asterisks indicate the AIY soma. The scale bar in (F) applies to (F’), and the one in (G) applies to (G’, H and H’). The length is 10μm. **(I)** Quantification of the fluorescent intensity of AIY RAB-3 for the indicated genotypes at the indicated stages. Data for each genotype are averaged from at least three biological replicates. The total number (N) of independent animals assayed is indicated in the bars. Statistical analyses are based on two-tailed Student’s t-test. *p<0.05, ***p<0.001, ****p<0.0001. Error bars represent SEM.

**Supplementary Fig. 7.**
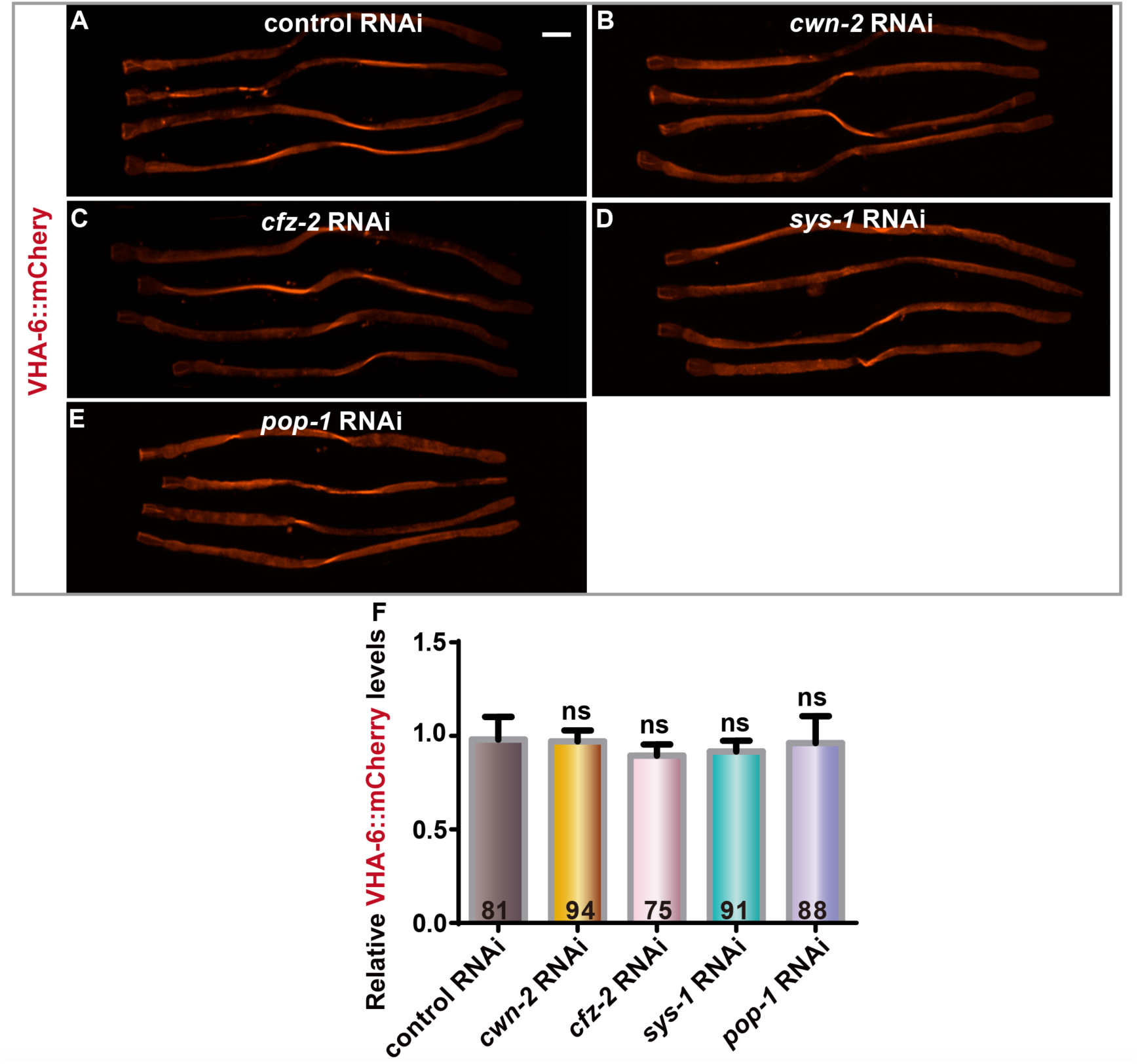
Wnt signaling does not affect VHA-6 expression. **(A-E)** Confocal images of an intestinal-specific reporter VHA-6::mCherry in control (A), *cwn-2 (*B), *cfz-2* (C), *sys-1* (D) and *pop-1* (E) RNAi animals at the adult day 1 stages. The scale bar in (A) is 50μm applies to all images. **(F)** Quantification of VHA-6::mCherry fluorescent intensity for the indicated genotype. Data are averaged from at least three biological replicates. The total number (N) of independent animals assayed is indicated in the bars. Statistical analyses are based on one-way ANOVA followed by Dunnett’s test. ns: not significant as compared to the control RNAi. Error bars represent SEM.

